# Benchmarking enrichment analysis methods with the disease pathway network

**DOI:** 10.1101/2023.09.29.560169

**Authors:** Davide Buzzao, Miguel Castresana-Aguirre, Dimitri Guala, Erik L.L. Sonnhammer

## Abstract

Enrichment analysis (EA) is a common approach to gain functional insights from genome-scale experiments. As a consequence, a large number of EA methods have been developed, yet it is unclear from previous studies which method is the best for a given dataset. The main issues with previous benchmarks include the complexity of correctly assigning true pathways to a test dataset, and lack of generality of the evaluation metrics, for which the rank of a single target pathway is commonly used.

We here provide a generalized EA benchmark and apply it to the most widely used EA methods, representing all four categories of current approaches. The benchmark employs a new set of 82 curated gene expression datasets from DNA microarray and RNA-Seq experiments for 26 diseases, of which only 13 are cancers. In order to address the shortcomings of the single target pathway approach and to enhance the sensitivity evaluation, we present the Disease Pathway Network, in which related KEGG pathways are linked. We introduce a novel approach to evaluate pathway EA by combining sensitivity and specificity to provide a balanced evaluation of EA methods. This approach identifies Network Enrichment Analysis methods as the overall top performers compared to overlap-based methods. By using randomized gene expression datasets, we explore the null hypothesis bias of each method, revealing that most of them produce skewed *p*-values.

## INTRODUCTION

Enrichment analysis (EA) is a popular approach for gaining biological insights from High-Throughput experiments. Gene expression profiles from DNA microarray or RNA-Seq research are commonly used as input data to EA. In a typical experiment, expression profiles for thousands of genes are obtained from a collection of samples from two categories such as case/control or treated/untreated, from which differentially expressed genes (DEGs) can be extracted. The goal of EA is to condense these profiles or DEG lists into a concise set of impacted biological functions or processes that can provide a systems level view of the changes and lead researchers to disease biomarkers or therapeutic targets. Functional gene sets representing molecular functions and biological processes are defined e.g. by the Gene Ontology (GO) [1], pathway databases such as Kyoto Encyclopedia of Genes and Genomes (KEGG) [2], Reactome [3], or experimentally derived gene set databases such as DisGeNET [4] or MSigDB [5].

EA methods can be grouped into four categories according to their procedural framework [6], that also mirrors the timeline of their development: (i) overlap-based methods test the significance of DEGs that overlap a functional gene set, (ii) per-gene scoring methods look for enriched ranking of a functional gene set in the sorted list of gene expressions, (iii) pathway topology methods acquire topological information from databases such as KEGG or Reactome to weigh the importance of each gene in the tested pathway, (iv) network-based methods look for enrichment of network links between DEGs and a functional gene set.

Despite the availability of a large range of EA methods, it is unclear which ones are the best for a given dataset. Various methods have been developed claiming to be better than DAVID [7] and GSEA [8], yet these remain highly cited methods for EA. However, many such benchmarks were based on simulated data. Tarca et al. [9] pioneered biological data benchmarking by using experimental datasets from diseases that could be associated with a particular pathway. Several recent enrichment evaluation studies have adopted this technique [10–13], yet the Tarca et al. benchmark has some problems such as only containing 19 diseases, 12 of which are cancer-related. Furthermore, to calculate sensitivity, the target pathway is treated as the single true positive and all other pathways are assumed to be false positives. The shortcoming of the single target pathway approach was first addressed by using mouse knockout experiments (KO) with a known KO gene, by considering all pathways that contain the KO gene as positive, and the rest as negative pathways [12]. However, this is only possible for a small number of datasets, and the validity of the positive and negative assignments can be questioned. Geistlinger et al. [13] updated the Tarca benchmark by including RNA-Seq data and a new scoring metric that uses MalaCards [14] to associate a disease dataset with more than one pathway. MalaCards scores disease relevance for a gene based on literature co-citation and experimental data, and summarizes per-gene relevance across the GO and KEGG gene sets. The benchmark calculates a “phenotype relevance” score based on the EA rankings and the precompiled relevance rankings. The approach is only based on relevance scoring and does not penalize false negatives, which is also the case for the Tarca benchmark. Because of this, the results tend to be biased to give an unfair advantage to methods that have low sensitivity but high specificity. Obtaining a good balance between these metrics is very important for a general benchmark.

The aim of this study is to present a generalized benchmark of the most popular EA methods, representing all four categories of existing approaches. As illustrated in Figure 1, the benchmark is structured in three parts. First, we provide a new collection of 76 curated DNA-microarray and 6 RNA-Seq gene expression datasets for 26 diseases, only 13 of which are neoplastic conditions. We ran 14 EA methods (two overlap-based, six per-gene scoring, four network-based and two topology-based) on the gene expression data against KEGG pathways. Second, we introduce the Disease Pathway Network to counteract the shortcoming of the single target pathway approach and improve the sensitivity assessment in an unbiased way. Third, we measure the sensitivity and specificity of the methods in an independent and balanced manner. Focusing on the performance under the null hypothesis, we further assess potential biases of each method and pathway.

**Figure 1.**
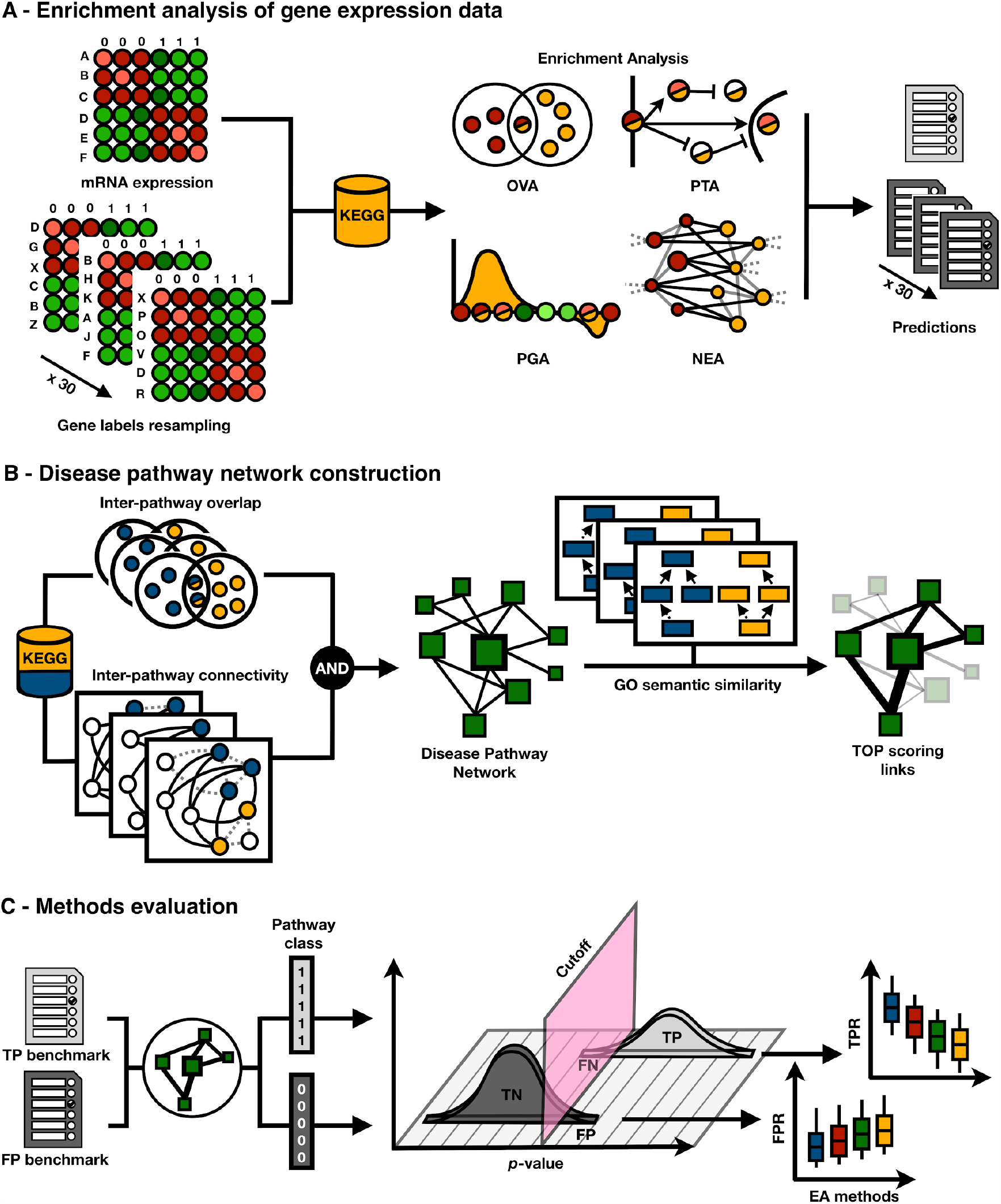
Procedural framework of the enrichment analysis benchmark. (A) EA methods are divided into four categories: OVerlap Analysis (OVA), Per-Gene score Analysis (PGA), Pathway Topology Analysis (PTA) and Network Enrichment Analysis (NEA), that differ in terms of the data and extra resources required for their input. All methods were run on 82 experimental expression datasets and on 2460 random datasets. (B) The KEGG pathway database is used to evaluate the EA methods. Each dataset is paired with the pathway that matches the disease and other pathways that are significantly related to the target pathway, as well as being top ranked by GO semantic similarity. Related pathways form the Disease Pathway Network which is based on both significant overlap and network connectivity. (C) Sensitivity (TPR) and specificity (1-FPR) are calculated for each method from their predictions on experimental and random datasets, respectively.

## MATERIALS AND METHODS

### Pathways database

KEGG is a popular collection of 16 databases that contain genomic information, biological pathways, disease- and drug-related data. We used the R package KEGGREST (v.1.34.0) to retrieve 318 KEGG (Release 101.0+/02-20, Feb 22) human pathways with at least 15 associated genes, that corresponds to the size of 95% of all the pathways. 92 of these pathways are related to human diseases, and belong to ten different subclasses i.e. Endocrine and metabolic disease, Neurodegenerative disease, Substance dependence, Infectious disease: bacterial, Infectious disease: parasitic, Infectious disease: viral, Cancer: overview, Cancer: specific types, Immune disease, Cardiovascular disease.

### Expression data

By querying the Gemma resource [15] using the 92 KEGG disease pathway names[15], we retrieved 82 datasets via the Gemma.R package (v0.99.1). Only datasets with a case-control observational study design, originally deposited in GEO [16], were included. Datasets were either of high quality during submission to GEO, or have undergone careful data and annotation curation in Gemma e.g. by flagging or removing outlier samples. In order to guarantee a high quality of gene expression, we required (i) a minimum number of three samples per condition (case or control), (ii) no drug treatment in the study, (iii) batch effect either not detected or corrected, (iv) a minimum of 15 differentially expressed genes at an FDR-corrected *p*-value of less than 0.2. DNA-microarray expression was quantile normalized and log-transformed. RNA-Seq experiments were retrieved as logarithm of counts per million reads (log_2_CPM). Probes were mapped to Gene IDs using the annotation table provided by Gemma, when multiple probes mapped to the same gene they were averaged. Basic analyses such as (co-)expression distribution plots, PCA analysis and differential expression analysis with volcano plots were performed to confirm the quality of each of the dataset, and reported as Supplementary Materials. For the “ulcerative colitis” datasets there was no exact match. Therefore, we used genes associated with “inflammatory bowel disease” of which “ulcerative colitis” is a subtype. A detailed description of each dataset is shown in Supplementary table 1.

All datasets were considered to have an unpaired design. limma (v3.54.0) was used to extract gene-wise moderated t-test scores of expression between case and control samples, run with trend=True for RNA-Seq datasets. Differentially expressed genes (DEGs) were defined as genes with Benjamini-Hochberg (BH) FDR-corrected *p*-values of less than 0.1 for 72 datasets, and of less than 0.2 for 10 datasets. When the number of DEGs exceeded 500, only the 500 most significant DEGs were selected.

### Disease pathway network construction

#### Inter-pathway connectivity

HumanNet-XC is a functional network of human genes that comprises inferred associations of gene co-expression, PPIs, genetic interactions, protein domain co-occurrence, and genomic context similarity from experimental datasets [17]. It was shown to have the highest protein coverage of all tested integrated human gene networks [17]. HumanNet shares gene ID nomenclature with KEGG meaning that its use also eliminated the need to accommodate the network’s gene vocabulary to pathways, which minimized inherent translation issues.

Inter-pathway connectivity *IPC(A,B)* was computed as the sum of the number of direct links *DL(A,B)* and shared neighbors *SN(A,B)* between the genes (e.g. *a* ∈ A, *b* ∈ B) of any two pathways (*e*.*g. A, B*), as connected in HumanNet-XC (v3) with 20% top scoring links (Eq. 1).

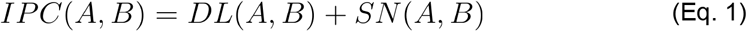

The number of direct links *DL(A,B)* was computed as the number of links between *A* and *B*.

The number of shared neighbors *SN(A,B)* was computed as the number of genes that were not part of pathways *A* and *B*, yet were connected to at least one gene in both *A* and *B*.

The *p*-value of the connectivity between two pathways was tested with a degree-aware subsampling test of 1,000 random samples, i.e. for the A against B test, A remained unchanged and B was replaced by a gene set where each gene was randomly sampled from genes with similar node degree in the network. This was achieved by sorting the 7248 KEGG genes by node degree and grouping them into bins of 100 genes. The binning made it possible to sample high-degree nodes that do not have a substitute with exactly the same degree. The *p*-value may be different when sampling A and B, for instance due to different sizes or degree distributions, hence we performed the test in both directions and combined the two *p*-values for each pathway pair using Fisher’s method [18].

#### Inter-pathway overlap

To calculate the significance of the gene overlap between two pathways, the same bi-directional degree-aware subsampling method as above was used, with 1,000 random samples. Here the overlap was expressed as Jaccard index *J(A,B)* (Eq. 2).

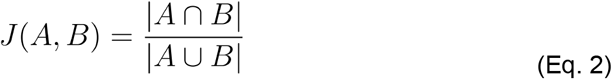

Only pathway pairs with FDR-corrected *p*-values of less than 0.05 from both inter-pathway connectivity and overlap tests were retained.

#### GO semantic similarity

For any two pathways that passed the overlap and network separation tests, the semantic similarity of Gene Ontology (GO) keywords was used as a proxy for their functional similarity. In particular, to compute similarity scores we used the Wang et al. graph-based method [19] which uses the topology of the GO Directed Acyclic Graph structures as implemented in the function clusterSim from the R package GOSemSim (v2.20.0). See Supplementary Materials for a more detailed description.

### Performance measures

In this study, the positive and negative benchmarks were independent. In the positive benchmark, true positives (TP) and false negatives (FN) were positive pathways (*i*.*e*. target or target-related) that were identified as significantly related (*p*-value<0.05), and not significantly related (*p-*value≥0.05), respectively. Similarly, in the negative benchmark, true negatives (TN) and false positives (FP) were negatives (*i*.*e*. unrelated pathways) that were identified as nonsignificant or significant, respectively. The negative benchmark was designed by resampling gene labels on the 82 datasets from the genome, for a total of 30x per dataset. Methods taking DEGs as input are inapplicable when only a few or nearly all genes are differentially expressed, but by utilizing gene label resampling we get the same number of DEGs per dataset as originally, and could estimate false positive rate (FPR) over 2460x318 tests for each method. To counteract the imbalance between positive and negative pathways, we considered only the number of target and target-related pathways from the positive benchmark to compute TN and FP in the negative benchmark. Having the definition of TP, TN, FP, and FN, we extracted true positive rate (TPR, or sensitivity) and true negative rate (TNR, or specificity, or 1-FPR) as follows:

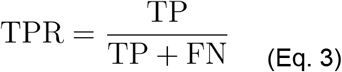

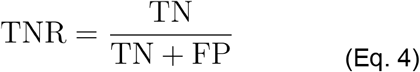

and used the geometric mean of TPR and TNR (G-mean) to condense the method performances in one balanced, comprehensive and robust summary index (Eq. 5).

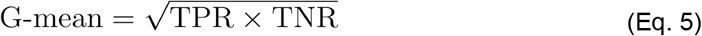

To underline the importance of collecting TPs among the top predictions, we also extracted the median relative rank of TPs in the predictions after sorting by *p-*value. In the presence of ties, we averaged the ranks.

### Enrichment Analysis methods

The EA methods selected for the benchmark are listed in Table 1. The methods under investigation belong to four different categories: (i) OVerlap Analysis (OVA), Per-Gene score Analysis (PGA), Pathway Topology Analysis (PTA) and Network Enrichment Analysis (NEA) methods. We summarize here the main characteristics of each category, see Supplementary Material for more details.

**Table 1.**
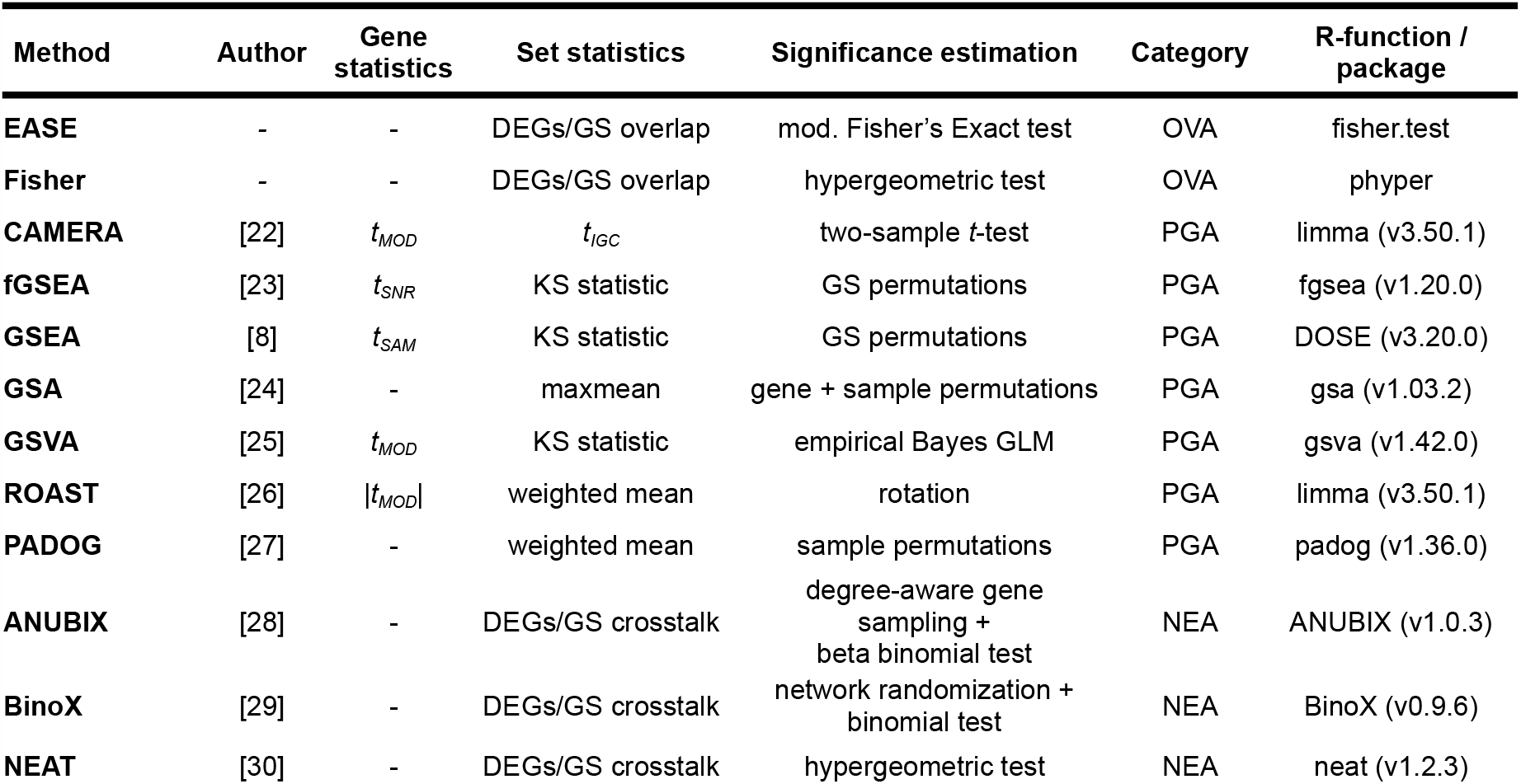
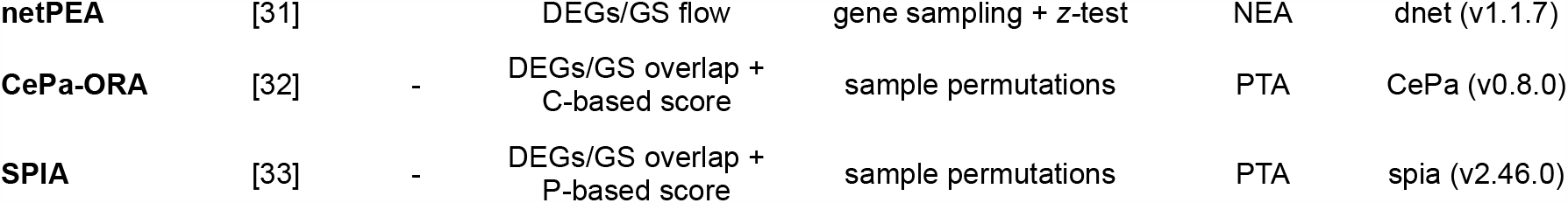
Enrichment Analysis methods under benchmark. OVerlap Analysis (OVA), Pathway Topology Analysis (PTA) and Network Enrichment Analysis (NEA) methods take Differentially Expressed Genes (DEGs) as input. Per Gene score Analysis (PGA) methods perform gene statistics via adaptations of Student’s *t*-test: *t*_*MOD*_ is moderated *t-*test, *t*_*SNR*_ is *t-*like signal-to-noise ratio, *t*_*SAM*_ is SAM’s *t* accounting for small variability at low expression values. All methods were run in an R (v4.1.0) version-controlled conda environment using the original packages, with the exception of EASE, fisher and netPEA that were run using our own R implementations, and BinoX that is a C++ program.

OVA methods test the proportion of DEGs in a functional gene collection against a discrete probability distribution model followed by extraction of one-sided *p*-values. PGA methods work with each gene separately, employing a statistical model to connect the response to the expression of each gene. Each gene undergoes a local statistical test that is used to determine a parametric or permutation-based *p*-value of variation of expression. A global statistical analysis is then performed to assign an enrichment score to a functional set of genes. For permutation-based methods, the significance of the global statistics for each functional gene set is then evaluated using 2000 sample permutations. Other settings are kept to default values, with the exception of CAMERA_fix that was run in two configurations: (i) CAMERA accounting for fixed inter-gene correlation of 0.01, (ii) CAMERA_flex extracting dataset-specific inter-gene correlation. PTA methods acquire topological information from databases such as KEGG or Reactome to weigh the importance of each gene in the tested pathway. PTA methods extract a per pathway score and perform size-aware permutations to extract *p-*values of enrichment. NEA methods evaluate the interconnectivity between DEGs and functional gene sets in the context of a functional association network, where a parametric or permutation-based approach is used to assign a *p*-value to the measured interconnectivity. FunCoup [20] and STRING [21] were used as underlying networks with a default confidence link score of 0.8 and 800, respectively.

## RESULTS

### The Disease Pathway Network for generalized benchmarking

To achieve high sensitivity in the benchmark we created the Disease Pathway Network, that connects target pathways to other related pathways. It was produced by first extracting 92 pathways from KEGG that are associated with human diseases, and connecting these to other KEGG pathways based on significant gene overlap and network connectivity. To rank these pathway-pathway relations in a functionally relevant way, we scored them using GO semantic similarity as described in the Supplementary Materials. To ensure an equal representation of the benchmarked disease pathway subnetworks, we restricted them to the top 20 linked pathways for each target disease pathway. This decision was motivated by the limited number of total associations observed in Parkinson’s and Huntington’s diseases (Supplementary Table 2). This resulted in a Disease Pathway Network of 212 pathways and 1520 links in total. Figure 2 shows a subnetwork of the target disease pathways in the Disease Pathway Network, with 143 pathways and 462 links. It illustrates the biological coherence of our disease pathway model, with stronger connections observed between diseases within the same class. Some pathways are very central in the Disease Pathway Network, with “Hepatitis C” having the largest number of connections. Our dataset is linked to 26 target diseases, and because disease representation varies, certain pathways were tested more often with our 82 benchmark datasets (Supplementary Figure 1). For example, the “Neurotrophin signaling pathway” was tested 61% of the time, while other pathways were only tested once.

**Figure 2.**
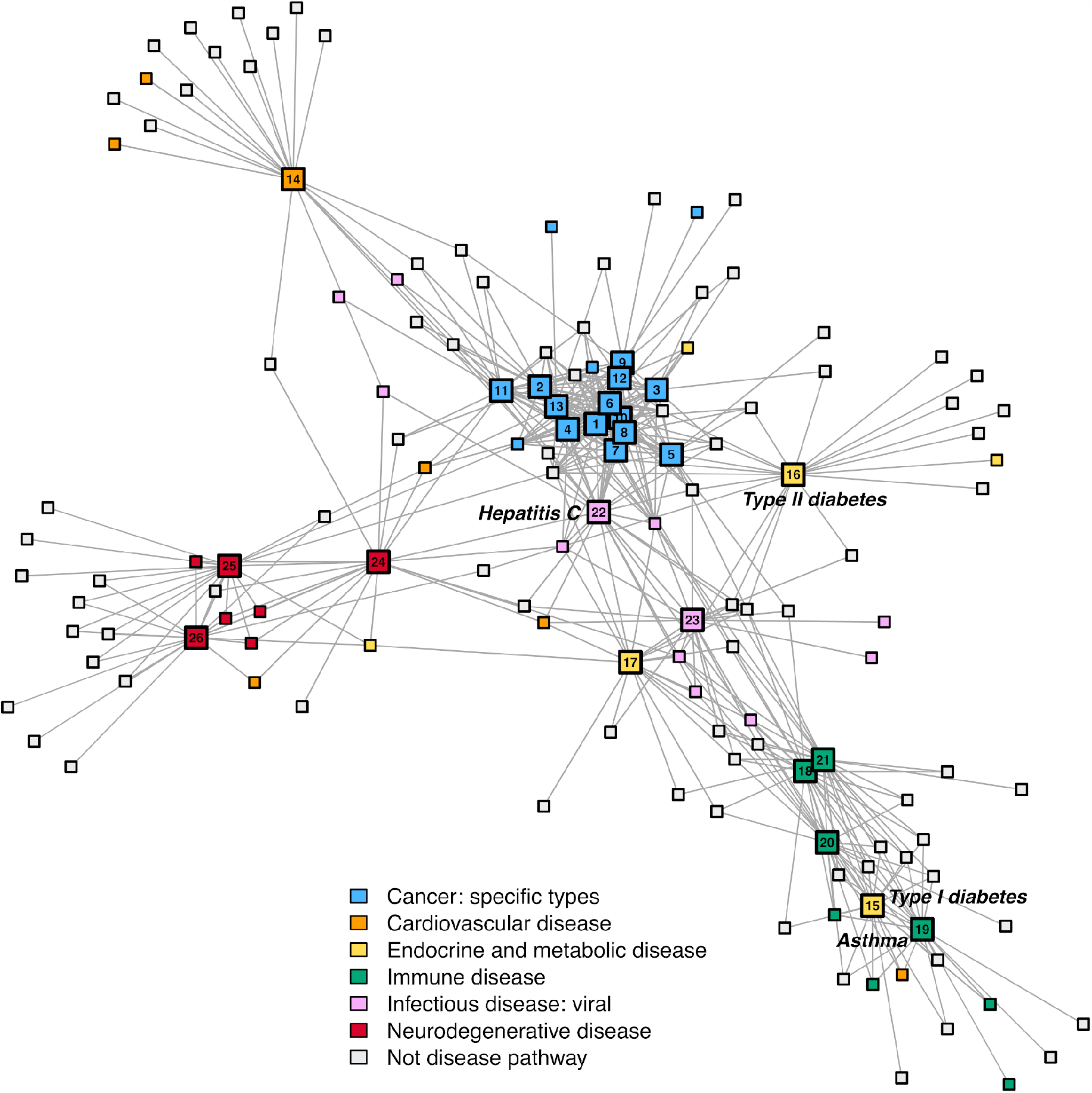
KEGG diseases and subclasses in the benchmark. An overview of the Disease Pathway Network, highlighting KEGG diseases and subclasses included in the benchmark. Target disease pathways are depicted with a larger node size and are labeled with an internal node id. See Supplementary Table 2 to match the pathway name and the node id. Links between non-target pathways are not shown as they are out of the scope of this benchmark. The colors correspond to the KEGG disease subclass a pathway belongs to. The width of the links represents the GO semantic similarity, which ranges from 0.77 to 0.98 in this network.

We also calculated the Jaccard index between all disease pathways within the same KEGG subclass under “Human Diseases” using the top 20 linked pathways (Supplementary Table 3). Here we observed that the “Cancer: specific types” subclass of pathways represents a highly connected subnetwork, with an average Jaccard index of 0.45, ranging from a minimum of 0.18 (“Glioma” - “Small cell lung cancer”) to a maximum of 0.82 (“Colorectal cancer” - “Pancreatic cancer”). On the other hand, some pathway subclasses were not interconnected, such as “Endocrine and metabolic disease” that includes type I and II diabetes. However, from a molecular mechanistic perspective it makes more sense that type I diabetes, an autoimmune disease [34,35], is connected to other autoimmune diseases such as Asthma rather than to type II diabetes. Another example of clustering across different disease classes is the “Hepatitis C” pathway, under “Infectious disease: viral”, that instead shows strong associations with cancers. This may be explained by the fact that “Hepatitis C” is a well-established risk factor for liver cirrhosis and liver cancer, despite primarily being a viral infection [36,37].

### Benchmarking 14 EA methods of different types

We then set up a benchmark where the Disease Pathway Network was used to identify target-related pathways for each gene expression dataset based on its target pathway. Fourteen EA methods of all four categories of currently existing EA approaches were benchmarked by considering also the target-related pathways as true positives. The methods were compared in terms of relative pathway ranking, sensitivity and specificity. To avoid introducing a bias towards a particular functional association network, we ran the NEA methods with two different networks, and refer to them with *-FunCoup and *-STRING in the method name. The performance of the investigated methods were summarized as a combination of average relative rank and G-mean of TPR and TNR (1-FPR) using the top 20 linked pathways per disease pathway in the Disease Pathway Network (Figure 3). Supplementary Table 4 provides a comprehensive breakdown of scores, including the distinctions between True Positives (TP), True Negatives (TN), False Positives (FP), and False Negatives (FN).

**Figure 3.**
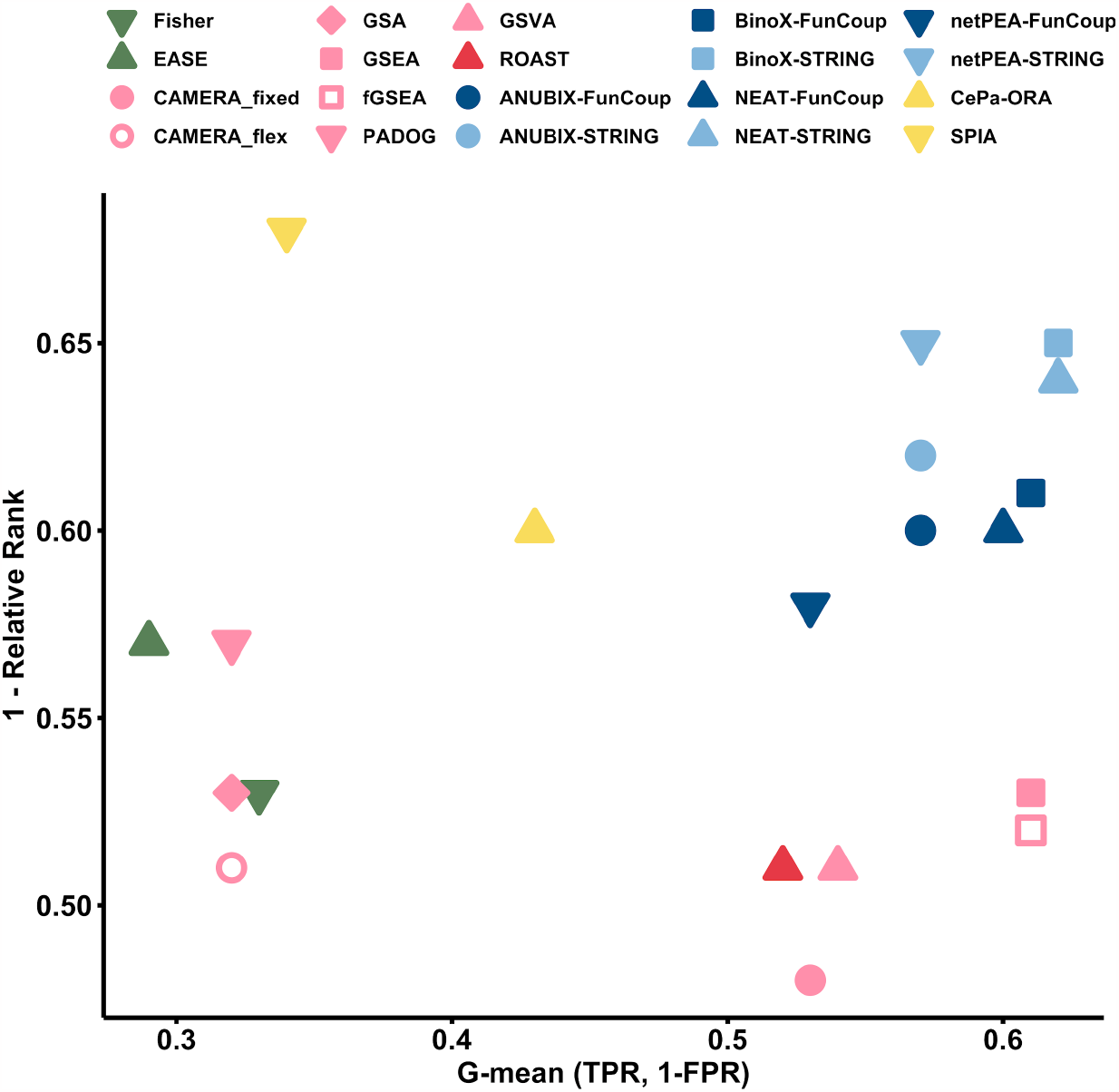
Summary of performance of 14 EA methods. G-mean of TPR and TNR (1-FPR) is shown versus 1 - relative rank, using the top 20 linked pathways per target pathway in the Disease Pathway Network. An ideal method would rank the related pathways for a specific dataset at the top of enrichments and would assign significant *p*-values to them. In the figure, colors refer to EA categories: green for OVA, pink for PGA with light and dark shades for competitive and self-contained null hypothesis, yellow for PTA, and blue for NEA methods with light and dark shades for STRING-/FunCoup-based implementations.

The NEA methods exhibited the best combined performance, with BinoX and NEAT reaching the highest G-mean at 0.62, and BinoX scoring 0.65 at 1-Relative Rank. The PGA and PTA methods performed well in only one of the two scoring metrics. Among the PGA methods, fGSEA and GSEA had best G-mean at 0.61, while PADOG had the best ranking at 0.57. Although the PTA method SPIA achieved the highest rank performance of all methods, it is important to note that this was inflated by the limited number of pathways that PTA methods can test (see details about PTA methods in Supplementary Material, Supplementary Figure 2). In our analysis, we assigned a *p*-value of 1 to all missing tests and computed the average rank for ties. A corollary of this is a low true positive rate, resulting in a poor G-mean. We noted that EASE got the lowest G-mean of 0.29, while CAMERA_fixed got the lowest rank performance at 0.48. A summary of the performance results with other cutoffs for the number of target-related pathways is shown in Supplementary Figure 3.

### Analysis of biases in the benchmarked methods

We also evaluated the performance of the EA methods on randomized data (Figure 4). An ideal method would generate a uniform distribution of *p-*values from 0 to 1 across pathways when randomized data is used, with 5% of the *p-*values being lower than the cutoff at 0.05. Five of the methods were extremely conservative, with a median FPR below 3%, and an FPR below the cutoff for all pathways. These include the OVA methods EASE and Fisher, the two CAMERA (PGA) methods, and the PTA method SPIA. The other PTA method, CePA-ORA, was however non-conservative with a median FPR of 9%.

**Figure 4.**
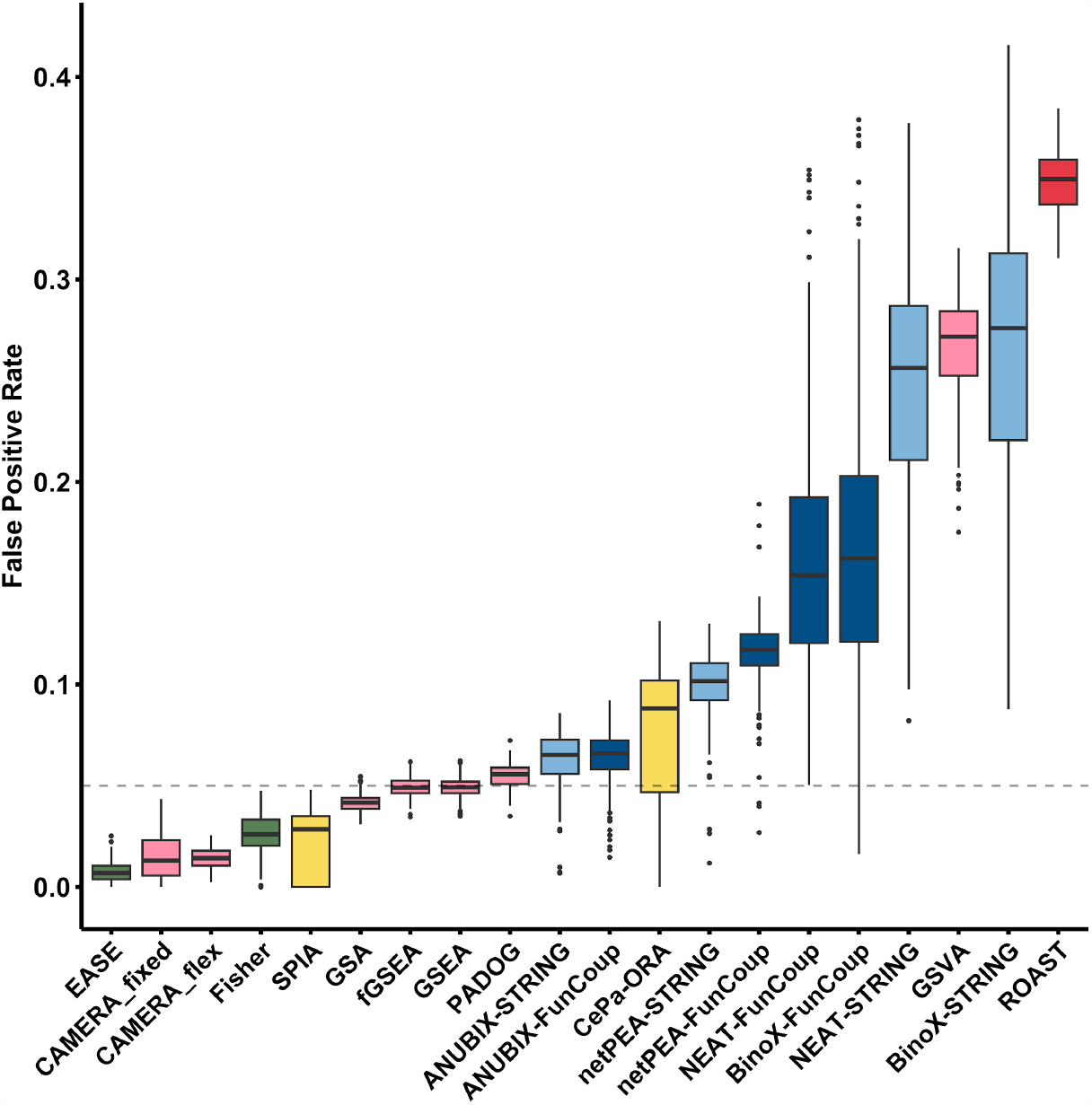
False positive benchmark. The false positive rates of 318 KEGG pathways for all benchmarked EA methods. The data was generated by resampling from the genome to generate random gene labels for the positive benchmark datasets. The gray dashed line represents the requested significance level of 0.05. The box colors correspond to method categories, as detailed in the caption of Figure 3.

At the other end, four methods were extremely non-conservative with a median FPR above 16% and and FPR above the cutoff for all pathways. These include the PGA methods ROAST and GSVA, and the NEA methods BinoX and NEAT. Both BinoX and NEAT were significantly more conservative when using FunCoup than STRING, with *p*=1.6e-53 and *p*=4.4e-55, respectively in a Wilcoxon rank sum test. In contrast, the FPR for netPEA was significantly higher when using FunCoup than STRING (*p*=1.9e-38). No significant difference between FunCoup and STRING was observed for ANUBIX (*p*=5.2e-1). The methods GSA, fGSEA, GSEA, PADOG and ANUBIX performed very well in this test with a median FPR close to 5%.

Under the null hypothesis, enrichment analysis methods often produce *p*-values that are either biased towards 0 or 1, or exhibit a bimodal distribution biased towards both extremes. Such a bias can affect the significance of the analysis, hence we examined the distribution of *p*-values for each method to see if it was right- or left-skewed (Figure 5). A right-skewed distribution (*p*-values biased towards 0) can potentially lead to false positives by reporting pathways as impacted when they are not. Conversely, a left-skewed distribution (*p*-values biased towards 1) can lead to false negatives by reporting pathways as not significant when they are actually impacted. GSEA was already reported to have no skewness bias [12] which we could confirm. Similarly, GSA, fGSEA and PADOG were unbiased in this benchmark. In contrast, the *p*-value distribution of the OVA methods EASE and Fisher was strongly skewed towards 1. At the other extreme, GSVA and ROAST had *p*-value distributions strongly skewed towards 0. BinoX, NEAT, and netPEA had distinct bimodal *p*-value distributions, mainly due to wrong assumptions about the EA score distribution, while ANUBIX showed almost no skewness. The bimodal *p*-value distribution of the PTA methods is due to the high number of untested pathways. For the tested pathways, SPIA had no bias, whereas CePA-ORA was righ-tskewed for all of them.

**Figure 5.**
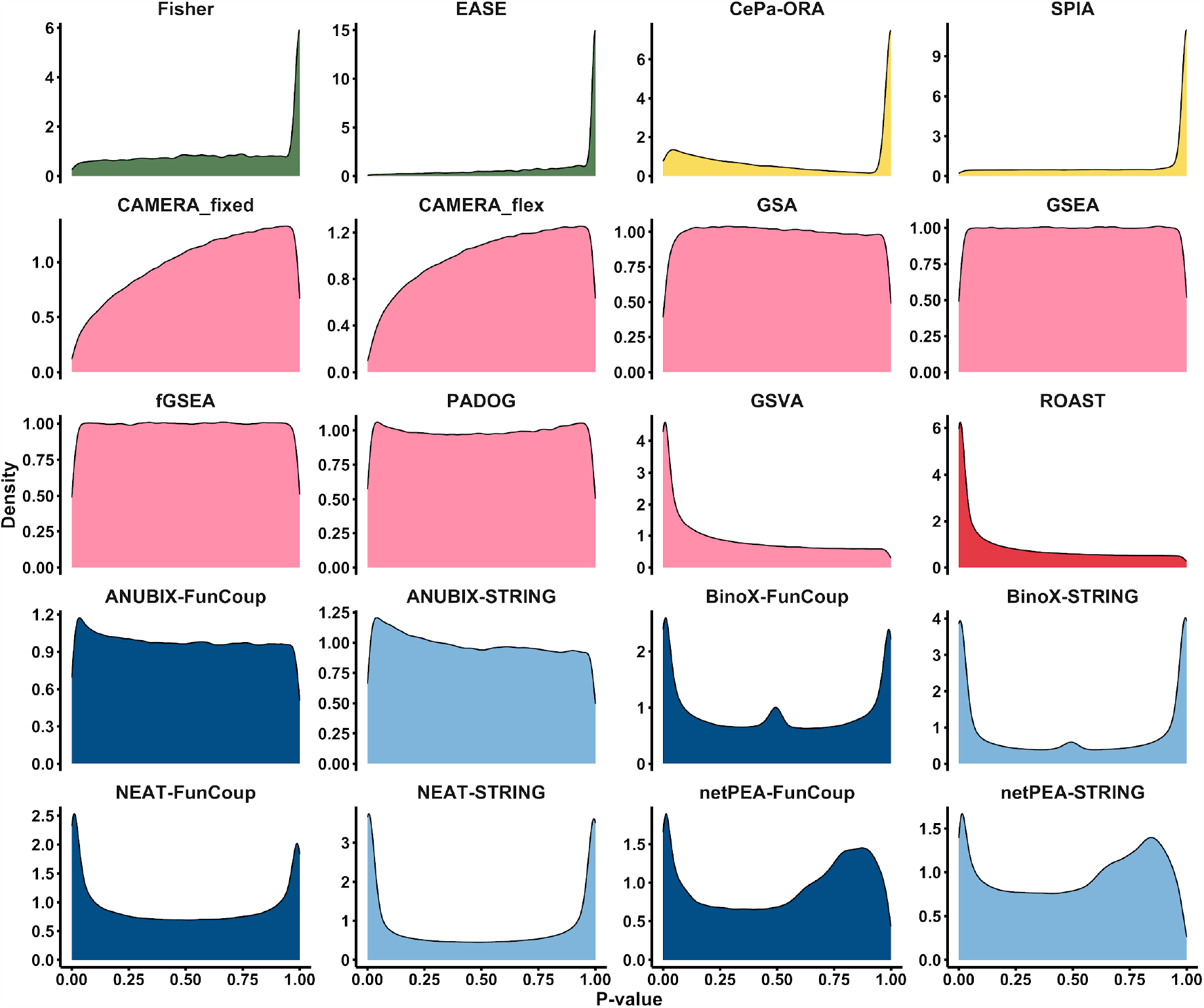
*P*-value distributions of all methods in the false positive benchmark. The *p*-values for all the pathways generated in the false positive benchmark are presented as density plots, depicting their distribution under the null hypothesis for each of the benchmarked methods. The ideal behavior is for the distribution to be uniform across the entire range from 0 to 1.

The NEA methods, which performed the best overall, still exhibited biases for some of the pathways that we decided to analyze further. As NEA methods rely on functional association networks, we expected the bias to be related to the underlying network topology. Thus, for each pathway under study, we calculated a number of network properties and then measured how they correlated with FPR (Supplementary Figures 4 and 5). We note a significant positive correlation for properties such as fraction of hubs, max degree, and size with both FunCoup and STRING. With FunCoup only, there was a strong correlation with fraction of intralinks, for the methods BinoX and NEAT, which has been previously noted [28]. A high fraction of intralinks means that the pathway is isolated and exhibits community-like properties in the network.

Further support is provided by a significant positive Spearman correlation of 0.66 for BinoX and 0.71 for NEAT between the FPR and the fraction of intralinks when using FunCoup (Supplementary Figure 5). BinoX and NEAT generalize the statistical properties of crosstalk for all pathways and are unable to adjust to unique properties of individual pathways, which are often highly non-random. In contrast, the correlation is weak in ANUBIX (0.06), which can be attributed to the fact that ANUBIX accurately predicts the expected degree of crosstalk by maintaining the pathways intact while performing a degree-aware resampling of the differentially expressed genes. No correlation was observed for netPEA that also does not alter pathway properties when constructing the null model. However, the correlation with the fraction of intralinks was much weaker when using BinoX and NEAT with the STRING network, which has other properties. On the other hand, using STRING led to a stronger correlation between skewness and some other network properties, including density of hubs, max degree, and median degree.

To highlight the pathways most prone to false detection in NEA methods, we isolated the top 10 pathways with the highest average FPR, as depicted in Figure 6A. We then standardized the distributions of network properties to visually represent the extent to which each pathway deviates from the mean (Figure 6B).

**Figure 6.**
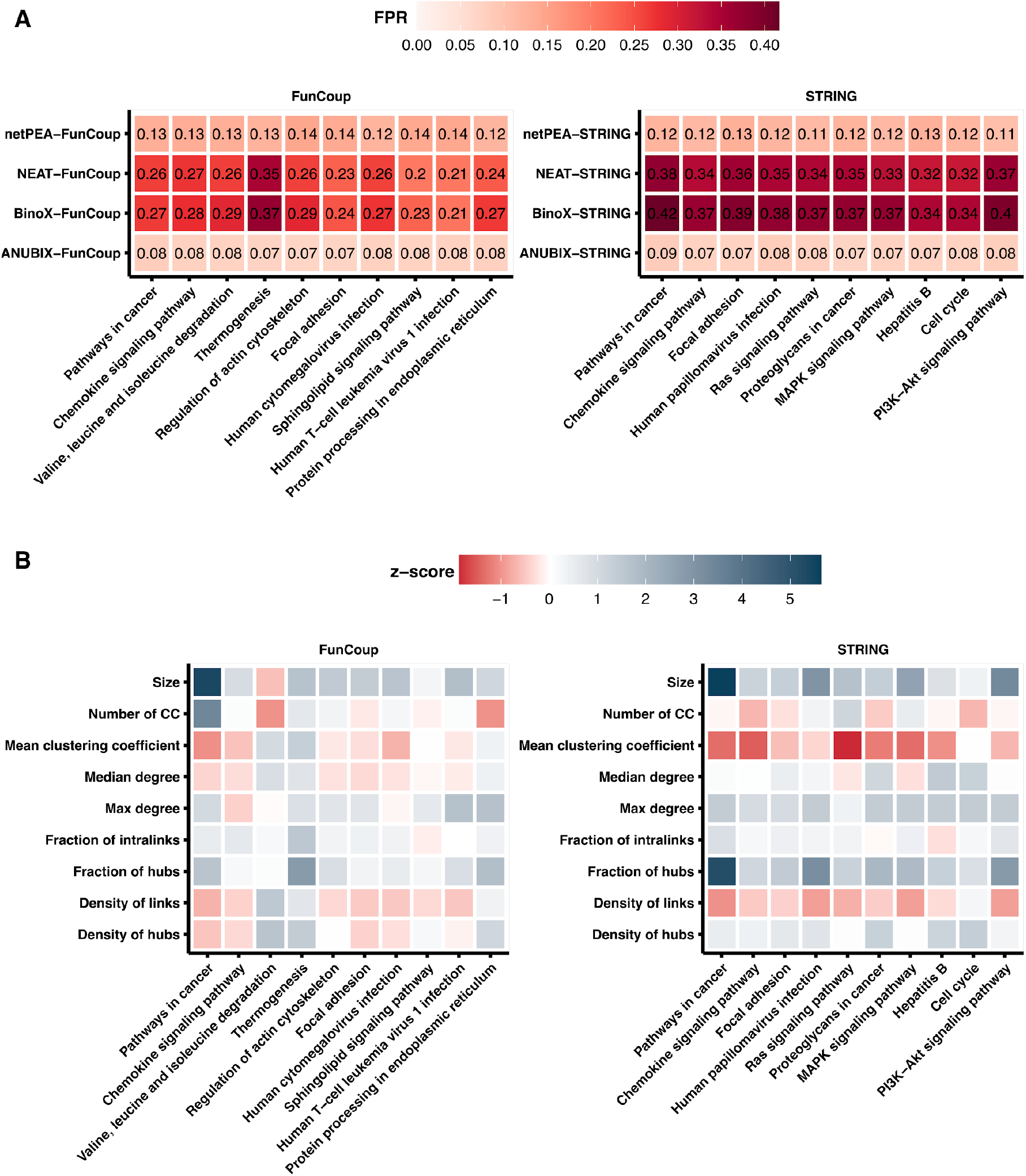
TOP falsely detected pathways in network-based EA methods. The heatmaps show (A) False Positive Rate (FPR) and (B) the z-score of the network properties of the 10 on average most falsely detected pathways predicted by the NEA methods, both for FunCoup and STRING. The FPR was computed per pathway over 2460 tests under the null hypothesis, and is the fraction of *p-*values < 0.05. See description of Supplementary Figure 4 for details on how the network properties were computed.

### Runtime evaluation

We performed scalability tests for each of the benchmarked methods. To detect trends on the runtime over all data in use, we show the runtime per dataset (Figure 7). The analysis was done on the 82 datasets of the benchmark with KEGG as input. GSA, GSEA, ROAST, and SPIA have internal parallelization. ANUBIX, BinoX, GSVA, NEAT, netPEA, and PADOG allow for parallelization and/or can treat multiple datasets at once. In a battery testing setup, this can vastly reduce the elapsed time. Network pre-processing in ANUBIX and BinoX, as well as KEGG pathways pre-processing in SPIA, was not included in the runtime because this only needs to be done once.

**Figure 7.**
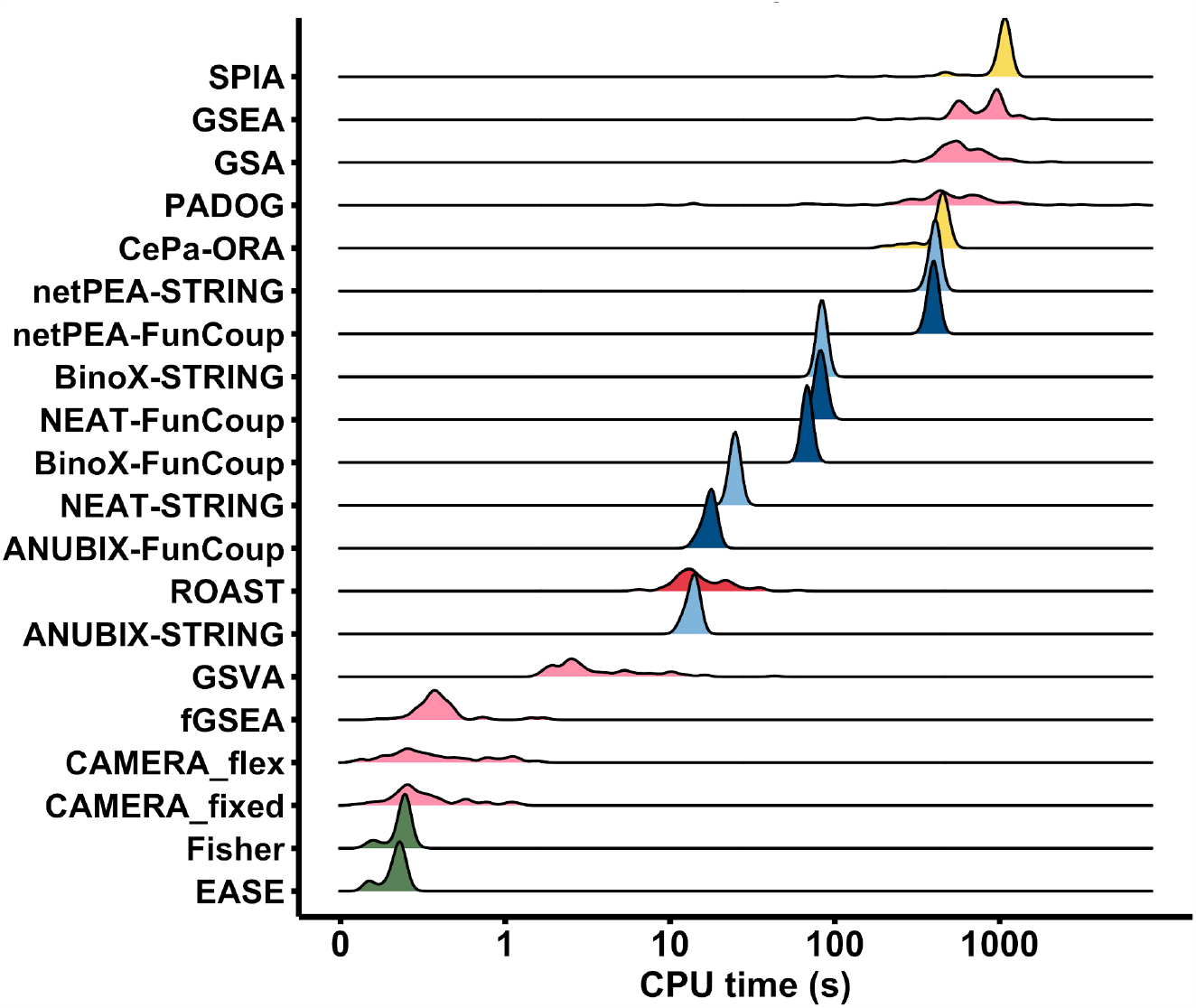
Runtime of enrichment analysis methods. Total CPU times when applying the enrichment methods to the 82 datasets of the benchmark. KEGG pathways were used to define gene sets. The computation was performed on an macOS Monterey (v.12.5.1) system with an Apple M1 processor (16GB RAM), with the exception of BinoX that was run on Ubuntu (v.18.04.6) system with Intel Core i7-2600 3.40GHz (16GB RAM). GSEA is shown as elapsed time.

## DISCUSSION AND CONCLUSION

We present a generalized benchmark for evaluating enrichment analysis (EA) methods on transcriptomic data that overcomes several limitations of previous benchmarks. This is the first benchmark that includes all four categories of EA methods, and it also covers a wider range of diseases than previously. While previous EA benchmarks have a strong bias towards cancer, we ensured that no more than half of our datasets were related to cancer. Our benchmark constitutes a further development of the performance assessment of EA methods by introducing the Disease Pathway Network based on connecting KEGG pathways, which overcomes the “single target pathway” shortcoming in a generalized way.

In contrast to previous benchmarks that separately evaluate sensitivity and specificity in pathway EA methods [9,12][13], our study uses an overall metric that combines both measures, providing a balanced evaluation of performance. This unified metric allows us to identify the best-performing method that excels in detecting enriched pathways while accurately avoiding non-enriched pathways. However, it is important to recognize the inherent difficulty in estimating the relative challenge of the False Positive (FP) benchmark compared to the True Positive (TP) benchmark. It appears that achieving a higher performance in the false positive (FP) benchmark is easier, but accurately quantifying the exact level of difficulty remains challenging. It is essential for future benchmarking efforts to carefully consider and communicate the trade-offs and challenges associated with both TP and FP benchmarks, providing a comprehensive understanding of method performance across different evaluation metrics.

Another drawback of previous benchmarks is that they were either aimed at methods that take gene sets as input [12,28], or at methods that use rankings of all genes. By catering for both types of methods, we were able to evaluate a total of 14 EA methods representing all four categories of currently existing approaches: (i) OVerlap Analysis (OVA), Per-Gene score Analysis (PGA), Pathway Topology Analysis (PTA) and Network Enrichment Analysis (NEA). The selected methods have a history of evaluation in prior benchmark studies [9,12,13,28]. All PGA methods tested in this benchmark are also implemented in EnrichmentBrowser [38]. To our knowledge, this is the first time that all four method categories have been evaluated using a standardized benchmark, providing valuable insights into the strengths and weaknesses of each category.

The benchmark showed that NEA methods have superior sensitivity compared to overlap-based methods, which may suffer from the limited coverage of knowledge-based databases like KEGG, GO, and Reactome [39]. Having a relatively high computational speed, NEA methods are well-suited for analyzing large-scale datasets efficiently. Among the NEA methods, BinoX and NEAT demonstrated the highest sensitivity at the price of high false positive rates, which appears to be related to bimodal *p*-value distributions. For BinoX and NEAT, this was partially a result of our approach to treat highly depleted pathways as non-significant enrichments. In addition to that, for pathways yielding a high variance of enrichment scores, statistical tests such as the binomial test (BinoX), the hypergeometric test (NEAT), and the Z-test (netPEA) are prone to underestimate the variance of the null distribution, resulting in a high false positive rate. ANUBIX, which employs the beta-binomial distribution to model overdispersed crosstalk distributions (see Supplementary methods), does not suffer from this and showed almost no skewness, achieving a good balance between sensitivity and specificity. Both netPEA and ANUBIX preserve pathway properties in their null model construction, resulting in absence of correlation between false positive rate and pathway fraction of intralinks in FunCoup. However, to account for biases in the underlying network, it is recommended to consider other factors than the fraction of intralinks, such as hub density and pathway size. In such a way, researchers can heed high false positive rates and enhance the reliability of their pathway EA. For users without programming experience, we can suggest user-friendly, web-based applications such as EviNet [40], PathwAX [41] and PathBIX [42] for a streamlined pathway NEA.

Recommendations for pathway analysis users can be tailored based on their specific research objectives. It is essential to highlight that OVA is an extremely conservative approach. The Fisher test, or its modified version, EASE, which is implemented in DAVID [7], although widely used for EA, has shown poor performance in the context of pathway analysis [29]. This can be attributed to the limited gene coverage of pathway databases and the assumption of gene independence, which does not hold true for genes interacting within pathways. As a result, users should exercise caution when relying on platforms using the Fisher test, such as Ingenuity Pathway Analysis [43] or DAVID, as they may produce false negatives due to the aforementioned limitations. Within PGA methods, we could support the ability of PADOG to be best at ranking the target pathways. CAMERA, despite its popularity, performed poorly in our benchmark, with worst results in the flexible configuration, whereas fGSEA/GSEA showed the best G-mean of sensitivity and specificity, with fGSEA being much faster. It can be conveniently executed using clusterProfiler [44], a comprehensive R package for gene set EA. PGA methods are valuable for predicting the direction and impact of biological function alterations. They uniquely consider gene expression changes, offering insights into pathway dynamics. For simulating the effects of expression changes, PTA methods like SPIA are the best choice. However, a drawback is that pathway topology is not always available, limiting their applicability. On the other hand, if researchers need to analyze arbitrarily defined sets of genes rather than predefined pathways, NEA methods may be more suitable provided that a functional association network is available for the species of interest. NEA methods differ from other approaches by considering the connectivity of differentially expressed genes with functional gene sets in a network context. Instead of assuming gene independence, NEA methods leverage genome-wide networks, such as FunCoup [20] and STRING [21], to capture the intricate interactions among genes within pathways. This network-focused analysis allows NEA methods to overcome the limitation of low database coverage leading to substantially higher sensitivity and an increased chance for obtaining biological insights.

In summary, choosing the appropriate pathway analysis method depends on the research objectives and the nature of the gene sets being analyzed. Researchers should carefully consider the benefits and limitations of EA methods to make informed decisions for their specific study designs. The limitations highlighted above collectively underscore the need for ongoing benchmarking efforts in the field of pathway EA. Ideally, a web-based platform would be developed to facilitate benchmarking of novel EA methods in a standardized and generalized way.

## DATA AVAILABILITY

We used R (r-project.org) v4.1.0 in a version-controlled conda environment for statistical tests and data visualization. The gene expression datasets can be retrieved from the Gemma database (https://gemma.msl.ubc.ca/). The code to reproduce the results of this benchmark can be found in the repository https://bitbucket.org/sonnhammergroup/EAbenchmark.

## ACKNOWLEDGEMENTS

The computations were enabled by resources provided by the National Academic Infrastructure for Supercomputing in Sweden (NAISS) and the Swedish National Infrastructure for Computing (SNIC) at PDC Center for High Performance Computing partially funded by the Swedish Research Council through grant agreements no. 2022-06725 and no. 2018-05973. This work was supported by the Swedish Research Council [2019-04095]. Open access funding is provided by Stockholm University.

## REFERENCES

1. Gene Ontology Consortium. Gene Ontology Consortium: going forward. Nucleic Acids Res. 2015; 43:D1049–56

2. Kanehisa M, Goto S, Sato Y, et al. Data, information, knowledge and principle: back to metabolism in KEGG. Nucleic Acids Res. 2014; 42:D199–205

3. Croft D, O’Kelly G, Wu G, et al. Reactome: a database of reactions, pathways and biological processes. Nucleic Acids Research 2011; 39:D691–D697

4. Piñero J, Ramírez-Anguita JM, Saüch-Pitarch J, et al. The DisGeNET knowledge platform for disease genomics: 2019 update. Nucleic Acids Res. 2020; 48:D845–D855

5. Liberzon A, Subramanian A, Pinchback R, et al. Molecular signatures database (MSigDB) 3.0. Bioinformatics 2011; 27:1739–1740

6. Khatri P, Sirota M, Butte AJ. Ten years of pathway analysis: current approaches and outstanding challenges. PLoS Comput. Biol. 2012; 8:e1002375

7. Hosack DA, Dennis G Jr, Sherman BT, et al. Identifying biological themes within lists of genes with EASE. Genome Biol. 2003; 4:R70

8. Subramanian A, Tamayo P, Mootha VK, et al. Gene set enrichment analysis: a knowledge-based approach for interpreting genome-wide expression profiles. Proc. Natl. Acad. Sci. U. S. A. 2005; 102:15545–15550

9. Tarca AL, Bhatti G, Romero R. A comparison of gene set analysis methods in terms of sensitivity, prioritization and specificity. PLoS One 2013; 8:e79217

10. Bayerlová M, Jung K, Kramer F, et al. Comparative study on gene set and pathway topology-based enrichment methods. BMC Bioinformatics 2015; 16:334

11. Dong X, Hao Y, Wang X, et al. LEGO: a novel method for gene set over-representation analysis by incorporating network-based gene weights. Sci. Rep. 2016; 6:18871

12. Nguyen T-M, Shafi A, Nguyen T, et al. Identifying significantly impacted pathways: a. comprehensive review and assessment. Genome Biol. 2019; 20:203

13. Geistlinger L, Csaba G, Santarelli M, et al. Toward a gold standard for benchmarking gene set enrichment analysis. Briefings in Bioinformatics 2021; 22:545–556

14. Rappaport N, Twik M, Nativ N, et al. MalaCards: A Comprehensive Automatically-Mined Database of Human Diseases. Curr. Protoc. Bioinformatics 2014; 47:1.24.1–19

15. Lim N, Tesar S, Belmadani M, et al. Curation of over 10 000 transcriptomic studies to enable data reuse. Database 2021; 2021:

16. Barrett T, Wilhite SE, Ledoux P, et al. NCBI GEO: archive for functional genomics data sets--update. Nucleic Acids Res. 2013; 41:D991–5

17. Kim CY, Baek S, Cha J, et al. HumanNet v3: an improved database of human gene networks for disease research. Nucleic Acids Res. 2022; 50:D632–D639

18. Fisher SRA. Statistical Methods for Research Workers. 1925;

19. Wang JZ, D. Z, Payattakool R, et al. A new method to measure the semantic similarity of GO terms. Bioinformatics 2007; 23:1274–1281

20. Persson E, Castresana-Aguirre M, Buzzao D, et al. FunCoup 5: Functional Association Networks in All Domains of Life, Supporting Directed Links and Tissue-Specificity. J. Mol. Biol. 2021; 433:166835

21. Szklarczyk D, Gable AL, Lyon D, et al. STRING v11: protein-protein association networks with increased coverage, supporting functional discovery in genome-wide experimental datasets. Nucleic Acids Res. 2019; 47:D607–D613

22. Wu D, Smyth GK. Camera: a competitive gene set test accounting for inter-gene correlation. Nucleic Acids Res. 2012; 40:e133

23. Korotkevich G, Sukhov V, Budin N, et al. Fast gene set enrichment analysis. bioRxiv 2021;

24. Efron B, Tibshirani R. On testing the significance of sets of genes. Ann. Appl. Stat. 2007; 1:107–129

25. Hänzelmann S, Castelo R, Guinney J. GSVA: gene set variation analysis for microarray and RNA-seq data. BMC Bioinformatics 2013; 14:7

26. Wu D, Lim E, Vaillant F, et al. ROAST: rotation gene set tests for complex microarray experiments. Bioinformatics 2010; 26:2176–2182

27. Tarca AL, Draghici S, Bhatti G, et al. Down-weighting overlapping genes improves gene set analysis. BMC Bioinformatics 2012; 13:136

28. Castresana-Aguirre M, Sonnhammer ELL. Pathway-specific model estimation for improved pathway annotation by network crosstalk. Sci. Rep. 2020; 10:13585

29. Ogris C, Guala D, Helleday T, et al. A novel method for crosstalk analysis of biological networks: improving accuracy of pathway annotation. Nucleic Acids Res. 2017; 45:e8

30. Signorelli M, Vinciotti V, Wit EC. NEAT: an efficient network enrichment analysis test. BMC Bioinformatics 2016; 17:352

31. Liu L, Ruan J. Network-based Pathway Enrichment Analysis. Proceedings 2013; 218–221

32. Gu Z, Wang J. CePa: an R package for finding significant pathways weighted by multiple network centralities. Bioinformatics 2013; 29:658–660

33. Tarca AL, Draghici S, Khatri P, et al. A novel signaling pathway impact analysis. Bioinformatics 2009; 25:75–82

34. Knip M, Siljander H. Autoimmune mechanisms in type 1 diabetes. Autoimmun. Rev. 2008; 7:550–557

35. Notkins AL, Lernmark A. Autoimmune type 1 diabetes: resolved and unresolved issues. J. Clin. Invest. 2001; 108:1247–1252

36. Perz JF, Armstrong GL, Farrington LA, et al. The contributions of hepatitis B virus and hepatitis C virus infections to cirrhosis and primary liver cancer worldwide. J. Hepatol. 2006; 45:529–538

37. Levrero M. Viral hepatitis and liver cancer: the case of hepatitis C. Oncogene 2006; 25:3834–3847

38. Geistlinger L, Csaba G, Zimmer R. Bioconductor’s EnrichmentBrowser: seamless navigation through combined results of set-& network-based enrichment analysis. BMC Bioinformatics 2016; 17:45

39. Gable AL, Szklarczyk D, Lyon D, et al. Systematic assessment of pathway databases, based on a diverse collection of user-submitted experiments. Brief. Bioinform. 2022; 23:

40. Jeggari A, Alekseenko Z, Petrov I, et al. EviNet: a web platform for network enrichment analysis with flexible definition of gene sets. Nucleic Acids Res. 2018; 46:W163–W170

41. Ogris C, Castresana-Aguirre M, Sonnhammer ELL. PathwAX II: network-based pathway analysis with interactive visualization of network crosstalk. Bioinformatics 2022; 38:2659–2660

42. Castresana-Aguirre M, Persson E, Sonnhammer ELL. PathBIX-a web server for network-based pathway annotation with adaptive null models. Bioinform Adv 2021; 1:vbab010

43. Krämer A, Green J, Pollard J Jr, et al. Causal analysis approaches in Ingenuity Pathway Analysis. Bioinformatics 2014; 30:523–530

44. Wu T, Hu E, Xu S, et al. clusterProfiler 4.0: A universal enrichment tool for interpreting omics data. Innovation (Camb) 2021; 2:100141

