## Supplementary Materials for "Benchmarking enrichment analysis methods with the disease pathway network"

### MATERIALS AND METHODS

#### GO semantic similarity

For any two pathways that passed the overlap and network separation tests, the semantic similarity of Gene Ontology (GO) keywords was used as a proxy for their functional similarity. In particular, to compute similarity scores we used the Wang et al. graph-based method [1] which uses the topology of the GO Directed Acyclic Graph (DAG) structures as implemented in the function clusterSim from the R package GOSemSim (v2.20.0), with the default semantic contribution factors $w_e$ of 0.8 and 0.6 for “is-a” and “part-of” relations, respectively. The similarity between the GO term $A$ and $B$ is given by the formula,

$$sim_{Wang}(A,B) = \frac{\sum_{t \in T_A \cap T_B} S_A(t) + S_B(t)}{SV(A) + SV(B)} \quad \text{(Eq. 1)}$$

Where the S-value $S_A(t)$ (or $S_B(t)$) scores the closeness of the DAG describing the relations of $A$ (or $B$) and all its ancestors in GO, with the DAG of a common GO term $t$ between $A$ and $B$. The S-value of GO term $t$ related to term $A$ is:

$$\begin{cases} S_A(A) = 1 \; if \; t = A \\ S_A(t) = max\{w_e \times S_A(t') | t' \in children \; of \; (t)\} \; if \; t \neq A \end{cases} \quad \text{(Eq. 2)}$$

$sim_{Wang}$ is thus the average of maximum semantic similarities, normalized by the semantic value of GO term $A$ and $B$, each computed as follows:

$$SV(A) = \sum_{t \in T_A} S_A(t) \quad \text{(Eq. 3)}$$

Because a protein can be annotated by multiple GO terms, the output of the analysis may be a matrix of scores which stores the similarity between the combinations of GO terms annotated for every couple of genes. A combining strategy known as Best-Match Average (BMA) was thus applied to average all best similarities between any combination of genes from two pathways. The analysis was run over all three GO divisions (i.e. Biological Process, Molecular Function and Cellular Component) by excluding Inferred Electronic Annotations (IEA), as IEA are not manually reviewed by experts. An average of the three scores was used as a proxy for the relatedness of the target pathway to any other pathway under study. In such a way we could build a weighted Disease Pathway Network of all target pathways and assess the sensitivity of the EA methods at different levels of confidence, *i.e.* extracting the top 20 connected pathways.

### Enrichment Analysis methods

The methods under investigation belonged to four different categories: (i) OVerlap Analysis (OVA), Per-Gene score Analysis (PGA), Pathway Topology Analysis (PTA) and Network Enrichment Analysis (NEA) methods. We report here algorithmic details of each EA category and method in the benchmark. Based on the underlying null hypothesis, an EA method can also be classified as competitive or self-contained [2]. A self-contained method tests if genes in the set of interest are differentially expressed whereas a competitive method tests if they are as differentially expressed as the genes not in the set. For a self-contained test on a gene set, the differential expression of only one of its genes may be enough to reject the null hypothesis that none of the genes in the gene set are different, overall resulting in increased sensitivity. However, self-contained methods might not be helpful in a battery testing setup given the large number of significant reported results. As a consequence, most of the available methods are competitive.

#### OVerlap Analysis (OVA)

OVerlap Analysis (OVA) methods test the proportion of differentially expressed genes (DEGs) in a functional gene collection against a discrete probability distribution model. One-sided *p*-values are extracted with a hypergeometric test in Fisher as implemented in [3], and with a modified Fisher's test in EASE, as implemented in DAVID [4].

#### Per-Gene score Analysis (PGA)

The most popular approach for analyzing expression data works with each gene separately, employing a statistical model to connect the response to each gene's expression. Each gene undergoes a local statistical test that is used to determine a parametric or permutation-based *p*-value of variation of expression. A global statistical analysis is then implemented to assign an Enrichment Score (ES) to a functional set of genes. For permutation-based methods, the significance of the global statistics for each functional gene set is then evaluated using 2000 sample permutations.

Gene Set Enrichment Analysis (GSEA) uses Kolmogorov-Smirnov (KS) statistics to test if a uniform distribution can be inferred from the rankings of the *p*-values of the genes in a gene set [5]. To derive the ES, GSEA implements a running-sum statistic that is increased (or decreased) upon encountering a gene inside (or outside) the gene set meanwhile traversing down from the top of the sorted gene list. NES is the maximum deviation from zero (ES) normalized by the size of the gene set. Here fast GSEA (fGSEA) is used to conduct a pre-ranked GSEA [6]. A *p*-value is estimated by permuting the genes in a gene set, which leads to randomly assigned gene sets of the same size.

Gene Set Analysis (GSA) substitutes the "maxmean" statistic (i.e. the maximum mean gene scores between the up- or down-regulated component of a gene set) for the weighted sign KS statistic, and scales it by its mean and standard deviation to obtain the re-standardized version of the "maxmean" statistic [7]. GSA generates a null distribution for estimating false discovery rates by combining gene and sample permutations.

ROtAtion gene Set Test (ROAST) implements a self-contained test that evaluates the enrichment of a gene set using rotation, a Monte Carlo simulation technology, rather than permutation [8].

Gene Set Variation Analysis (GSVA) computes sample-wise gene set enrichment scores as a function of genes within and outside the gene set, transforming the data from a gene by sample matrix to a gene set by sample matrix [9]. With the goal of assessing pathway variation in large, diverse populations with complex phenotypic traits, GSVA assesses variance in gene set enrichment across samples regardless of class identification (*i.e.* with no assumption of case/control). As recommended in the manual, we used limma to examine variations in enrichment scores between sample groups.

Correlation Adjusted MEan RAnk gene set test (CAMERA) implements a test that accounts for gene-to-gene correlations [10]. CAMERA was run in two configurations: (i) CAMERA_fixed accounting for fixed inter-gene correlation of 0.01; (ii) CAMERA_flex extracting dataset-specific inter-gene correlation.

Pathway Analysis with Down-weighting of Overlapping Genes (PADOG) pre-computes a table with gene frequencies in the gene sets database and then extracts a ES for each gene set by assigning the mean of absolute values of moderated gene t-scores, weighted by the gene frequencies [11]. In this way, genes that are found in fewer gene sets have a great impact in the analysis, whereas the contribution of ubiquitous genes is down-weighted. The ES is tested against the null distribution, generated by sample permutations.

#### Pathway Topology Analysis (PTA)

Signaling Pathway Impact Analysis (SPIA) tests the abnormal perturbation of a given pathway on top of the over-representation of DEGs in the same pathway, as it works in the ORA method [12]. The perturbation of a pathway is determined by propagating fold-changes in expression across the pathway topology. The pathway topology can be represented with a signed directed graph, with the sign of the edges corresponding to the type of interaction. To generate the SPIA pathway input, we converted each pathway graph into two adjacency matrices, one for activation processes and the other for inhibition processes. To represent activation and inhibition, we assigned a weight vector β where 1 denotes activation and -1 denotes inhibition. To assess the significance of a pathway perturbation, SPIA compares the observed net perturbation against the perturbation computed in a randomized scenario in which as many genes as the DEGs are allowed to occupy any position along the pathway and to have any log fold-change within the range of DEGs values. A final *p*-value is returned as the combination of the two analyses, by default using the Fisher method. SPIA does not return a functional enrichment for roughly 1/3 of total pathways, as metabolic pathways in KEGG do not include any activation and/or repression interaction.

Centrality Pathway Over Representation Analysis (CePa-ORA) uses pathways as networks with nodes representing genes [13]. For each pathway, CePa-ORA measures node weights using various centrality metrics: in-degree, out-degree, betweenness, in-largest reach, out-largest reach, and equal weight condition. The final pathway score is calculated by summing the weights of nodes that resulted differentially expressed in the DE analysis. Pathway significance is assessed using a null distribution derived from permutation of DE genes. Similarly to *Nguyen et al. 2019* [14], we selected the final *p*-value for each pathway as the lowest *p*-value among the six derived from the different centrality measurements, as the authors did not indicate a preferred metric.

#### Network Enrichment Analysis (NEA)

NETwork-based Pathway Enrichment Analysis (netPEA) executes Random Walk with Restarts (RWR) on a network by using DEGs as seed nodes with a predetermined restart probability, by default of 0.5 [15]. At steady state, the resulting probabilities will contain the likelihood of RWR ending in each network node that may be used to derive a distance between the DEGs and all other nodes in the graph. NetPEA averages the probabilities between all genes in a gene set to obtain a final ES. z-scores are calculated to extract *p*-values by extracting a null distribution with a size-aware resampling procedure of size 2000.

Network Enrichment Analysis Test (NEAT) finds crosstalks between the DEGs and a gene set. The crosstalk is defined as the degree of interconnectivity between two gene sets mapped onto a functional association network. NEAT tests enrichment (or depletion) significance against a hypergeometric discrete null distribution of crosstalks [16]. Naïvely, it calculates the expected number of links between two gene sets based on their degrees and then uses the hypergeometric distribution to assess statistical significance. In this way the method assumes that under no functional enrichment the crosstalk follows a hypergeometric distribution, thus the analysis speeds up.

BINOmial null distribution of X-talk (BinoX) asserts the crosstalk between the DEGs and a gene set is statistically significant if it is not expected from a null model based on randomisation of the network [17]. By default, BinoX extracts N randomized networks based on the Link Assignment and Second Order Conservation method (LA+S), which preserves the original second order topological properties and the network degree distribution within the random model. Once the parameters of the random model have been determined, they may be used to assess the likelihood of observed crosstalk. Because all network connections are binary, it is assumed that crosstalk between any two gene sets follows a binomial distribution. As for NEAT and BinoX, if the expected crosstalk is less than the observed, the *p*-value for the upper tail represents enrichment, whereas the opposite may be interpreted as depletion and is calculated based on the lower tail of the distribution. As depletion is out of the scope of the benchmark, we take 1 - *p*-value as significance for all gene sets with smaller crosstalk than expected.

Adaptive NUll distriButIon of X-talk (ANUBIX) tests the crosstalk between the DEGs and a gene set [18]. Statistical significance is assessed by testing the crosstalk against a per DE gene set adapted beta-binomial distribution estimated by computing a discrete null distribution of crosstalk computed with a degree-aware resampling procedure of size 2000.

KEGG is chosen to compare methods throughout the analysis. To accommodate each network's gene vocabulary, *UP000005640_9606.idmapping* (https://www.uniprot.org/news/2021/02/10/release) and *9606.protein.aliases.v11.5.txt* (https://string-db.org/cgi/download?sessionId=bDexq6ZYgwXq&species_text=Homo+sapiens) are used to map *Gene ID* and *Gene Name* to *Ensembl* gene and protein identifiers, for FunCoup and STRING respectively.

### Supplementary Figures

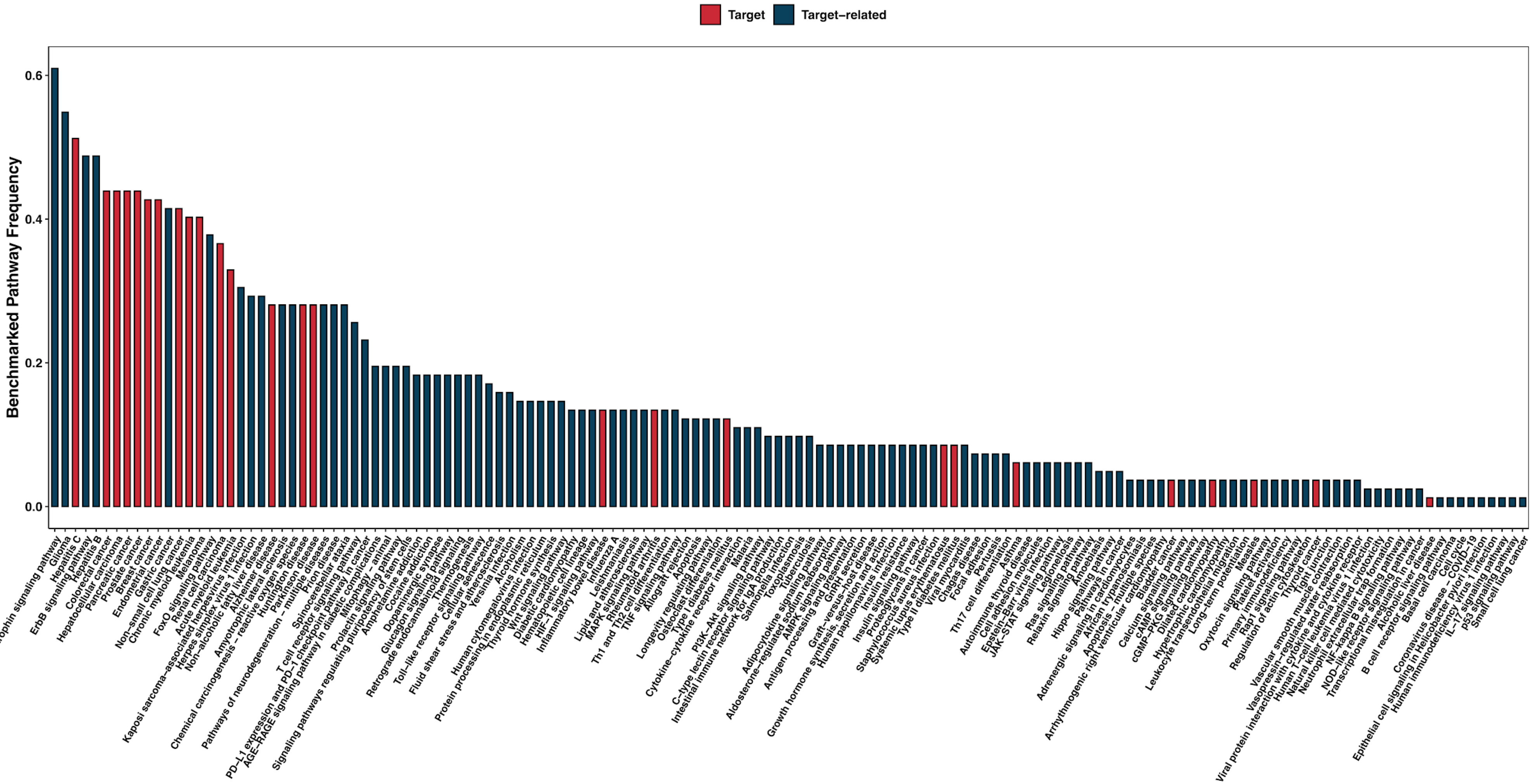


**Supplementary Figure 1 - Fraction of times a pathway is tested in the benchmark.**

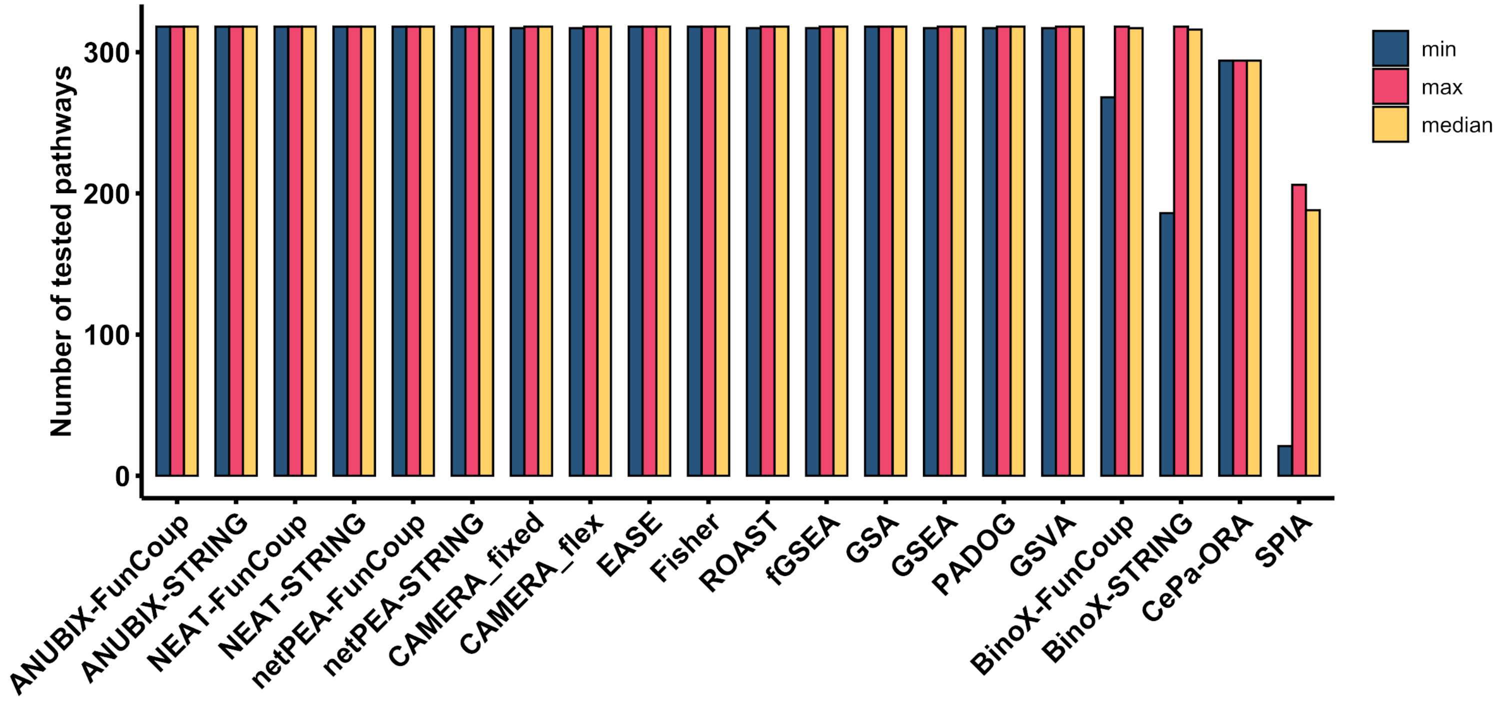


**Supplementary Figure 2 - Min, max and median average number of tested pathways in the positive benchmark.**

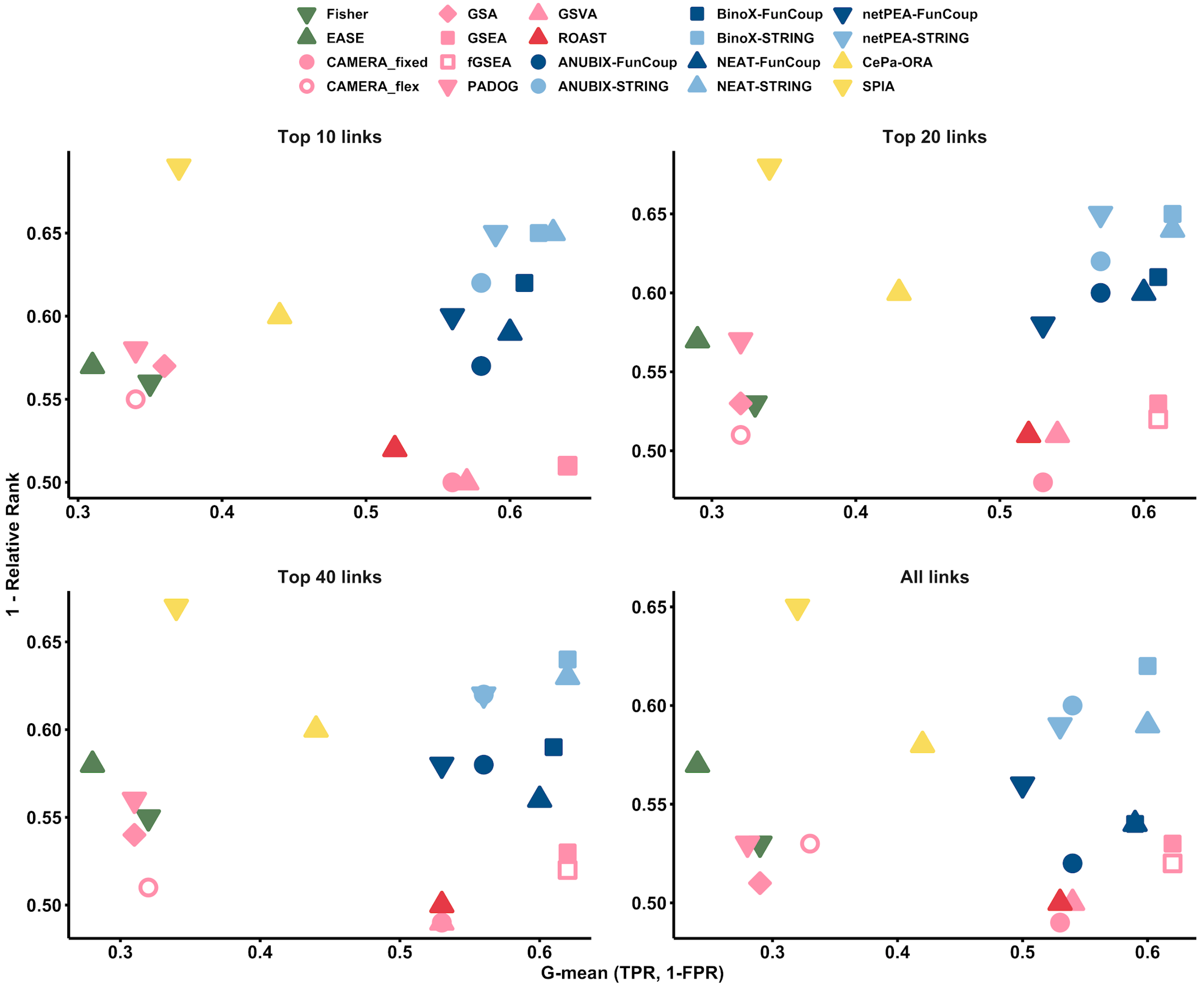


**Supplementary Figure 3 - Summary of performances as Relative Rank against G-mean of TPR and FPR.**

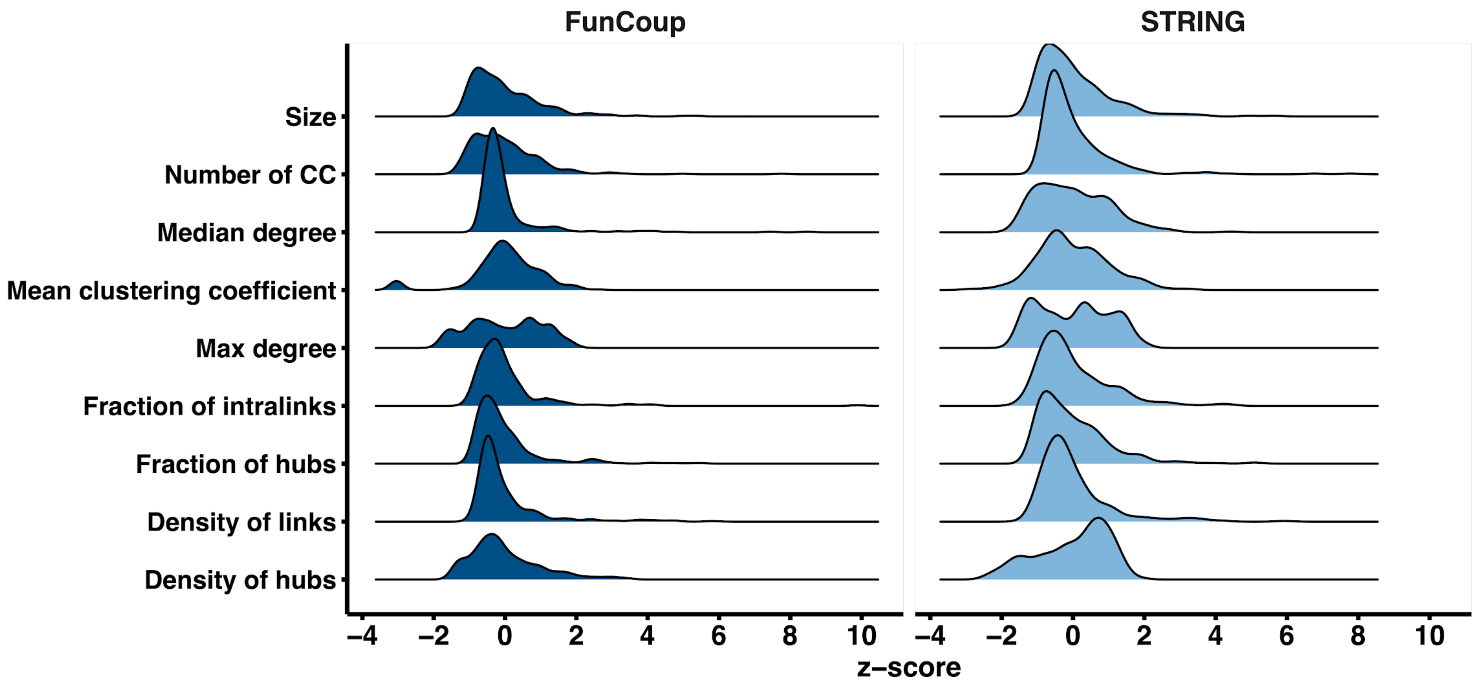


**Supplementary Figure 4 - Distribution of standardized network properties of KEGG pathways in FunCoup and STRING.** The pathway properties are: size of the pathway; the fraction of intralinks as the ratio of the number of links within the pathway and the overall number of links for the same pathway genes in the network; the density of links as the proportion of existing links within the pathway against the total number of possible links; the max and median average pathway degree; the mean average clustering coefficient; the number of connected components (CC); the fraction of hubs as the ratio of the number of hubs within the pathway and the overall number of pathway nodes; and the density of hubs as the proportion of existing hubs within the pathway against the total number of network hubs, which is 2411 in FunCoup and 3248 in STRING. We defined "hub" as any node with degree k when p(x>k) is smaller than 0.05 in a Poisson distribution with lambda equal to the network average degree, which was 95 and 25 for FunCoup and STRING, respectively.

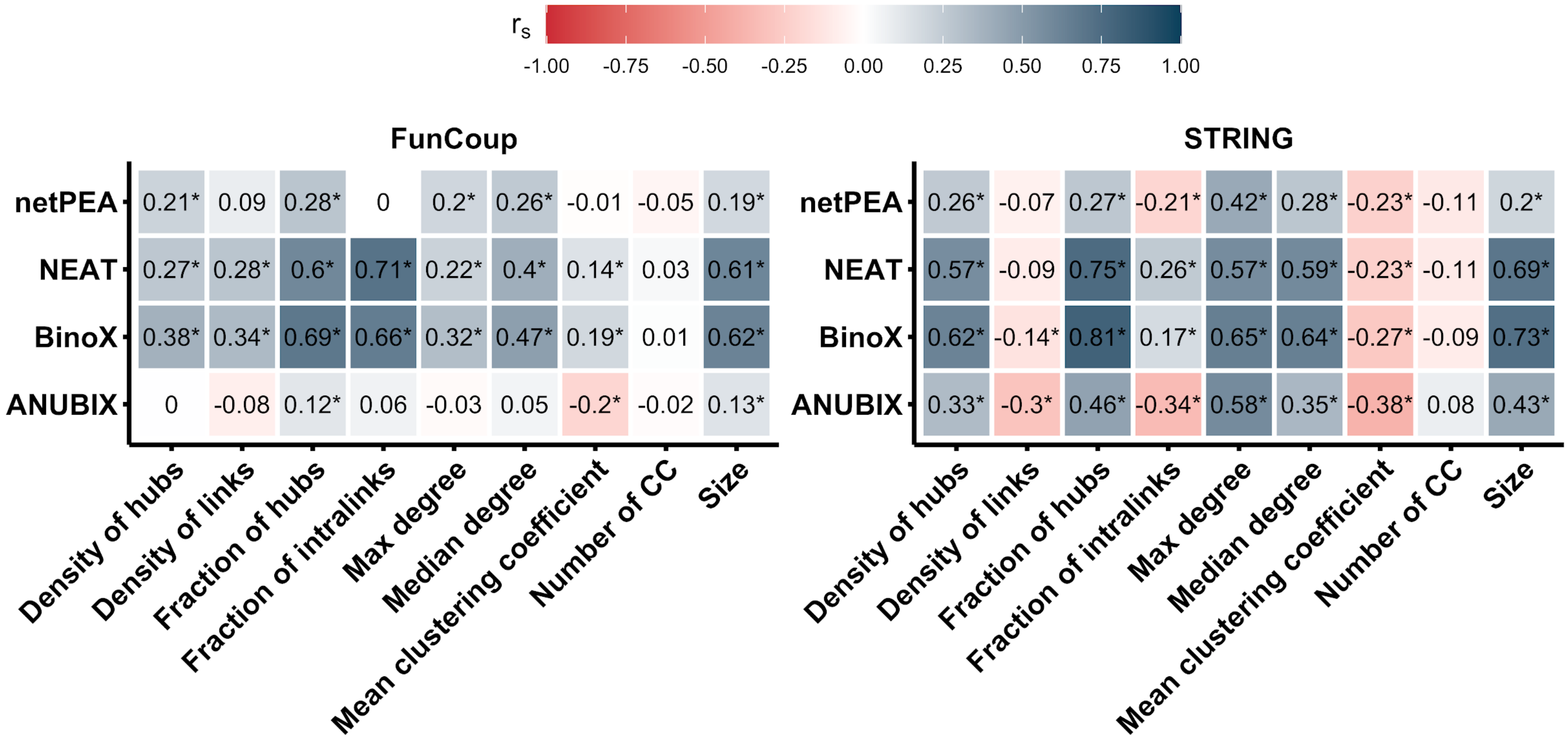


**Supplementary Figure 5 - Correlation between pathway network properties and false positive rate.** The heatmap shows Spearman correlations for a selection of network properties against the False Positive Rate (FPR) of KEGG pathways as predicted by network-based EA methods with FunCoup and STRING as underlying functional association networks. See description of Supplementary Figure 4 for details about computing the network properties. *P*-values of the correlations were estimated with a test for correlation between paired samples and are reported in Supplementary Table 5. Significant correlations with *p*-values below 0.05 are shown in the figure with a *. The FPR was evaluated on the benchmark datasets by resampling from the genome to generate random gene labels on the 82 datasets. This resampling process was repeated 30 times per dataset, resulting in a total of 2460 resampling tests per pathway. FPR was computed as the complement to 1 of TNR (eq. 7).

### Supplementary Tables

**Supplementary Table 1 - Overview of data.** 82 well-characterized DNA-microarray and RNA-Seq datasets in which a pathway is assumed to be prior knowledge under the assumption of an association with the tested condition were retrieved from GEMMA, 76 of which are DNA-microarray and 6 RNA-Seq (*). GEO accession is reported. The number of samples was chosen to be at least 3 per condition. Differentially Expressed Genes (DEGs) are genes with Benjamini-Hochberg (BH) FDR-corrected *p*-values of less than 0.1 and of less than 0.2 (†). The number of DEGs was set to be no larger than 500. The number of control and case samples, as well as up- and down-regulated genes, is shown within parentheses under respective columns in the table.

| GEO | Disease/Target pathway | Nr. of samples | Genome coverage | Nr. of DEGs | Batch effect |
|---|---|---:|---:|---:|---|
| **GSE14858** | Acute myeloid leukemia | 39 (20/19) | 18337 | 500 (293/207) | Corrected |
| **GSE28619** | Alcoholic liver disease | 22 (15/7) | 18464 | 500 (331/169) | Not detected |
| **GSE12685** | Alzheimer disease | 13 (6/7) | 11141 | 500 (222/278) | Corrected |
| **GSE28146** | Alzheimer disease | 29 (21/8) | 17912 | 500 (171/329) | Corrected |
| **GSE36980** | Alzheimer disease | 79 (32/47) | 17837 | 500 (141/359) | Corrected |
| **GSE37263†** | Alzheimer disease | 16 (8/8) | 15133 | 500 (169/331) | Corrected |
| **GSE39420** | Alzheimer disease | 21 (14/7) | 17947 | 500 (162/338) | Not detected |
| **GSE4757** | Alzheimer disease | 20 (10/10) | 18801 | 94 (65/29) | Corrected |
| **GSE95587*** | Alzheimer disease | 117 (84/33) | 18297 | 500 (211/289) | Not detected |
| **GSE97760** | Alzheimer disease | 19 (9/10) | 10779 | 500 (397/103) | Not detected |
| **GSE19187** | Asthma | 38 (27/11) | 17803 | 500 (366/134) | Corrected |
| **GSE23552** | Asthma | 39 (26/13) | 15210 | 500 (290/210) | Corrected |
| **GSE27011** | Asthma | 54 (36/18) | 17634 | 500 (273/227) | Corrected |
| **GSE31189** | Bladder cancer | 92 (52/40) | 18649 | 120 (50/70) | Corrected |
| **GSE10810** | Breast cancer | 58 (31/27) | 18543 | 500 (77/423) | Corrected |
| **GSE26304** | Breast cancer | 115 (109/6) | 16638 | 500 (107/393) | Corrected |
| **GSE43754** | Chronic myeloid leukemia | 19 (9/10) | 15020 | 500 (308/192) | Not detected |

| | | | | | |
|---|---|---|---|---|---|
| **GSE10715** | Colorectal cancer | 30 (19/11) | 18210 | 500 (244/256) | Corrected |
| **GSE13067** | Colorectal cancer | 72 (11/61) | 18722 | 500 (279/221) | Corrected |
| **GSE31737** | Colorectal cancer | 79 (40/39) | 15188 | 500 (292/208) | Corrected |
| **GSE4107** | Colorectal cancer | 22 (12/10) | 18618 | 500 (297/203) | Corrected |
| **GSE49355** | Colorectal cancer | 56 (39/17) | 11233 | 500 (192/308) | Corrected |
| **GSE50117** | Colorectal cancer | 18 (9/9) | 10100 | 500 (311/189) | Not detected |
| **GSE16499** | Dilated cardiomyopathy | 30 (15/15) | 15095 | 500 (218/282) | Corrected |
| **GSE3586** | Dilated cardiomyopathy | 28 (13/15) | 3765 | 500 (231/269) | Not detected |
| **GSE42955** | Dilated cardiomyopathy | 29 (24/5) | 17854 | 500 (169/331) | Corrected |
| **GSE36389** | Endometrial cancer | 19 (13/6) | 11210 | 500 (151/349) | Corrected |
| **GSE40184** | Hepatitis C | 18 (10/8) | 11099 | 500 (240/260) | Corrected |
| **GSE38476** | Hepatocellular carcinoma | 20 (10/10) | 13131 | 500 (288/212) | Not detected |
| **GSE54236** | Hepatocellular carcinoma | 160 (80/80) | 16823 | 500 (300/200) | Corrected |
| **GSE45516** | Huntington disease | 9 (6/3) | 18331 | 500 (339/161) | Not detected |
| **GSE64810*** | Huntington disease | 69 (20/49) | 16046 | 500 (338/162) | Not detected |
| **GSE73655** | Huntington disease | 20 (13/7) | 19881 | 65 (39/26) | Not detected |
| **GSE14580** | Inflammatory bowel disease | 14 (8/6) | 18612 | 500 (273/227) | Corrected |
| **GSE22619** | Inflammatory bowel disease | 20 (10/10) | 18819 | 500 (201/299) | Corrected |
| **GSE36807** | Inflammatory bowel disease | 35 (28/7) | 18554 | 500 (193/307) | Not detected |
| **GSE5808** | Measles | 18 (15/3) | 11156 | 500 (135/365) | Corrected |
| **GSE15605** | Melanoma | 74 (58/16) | 18045 | 500 (145/355) | Not detected |
| **GSE18842** | Non-small cell lung cancer | 91 (46/45) | 18683 | 500 (205/295) | Corrected |
| **GSE19188** | Non-small cell lung cancer | 156 (91/65) | 18484 | 500 (141/359) | Corrected |
| **GSE19804** | Non-small cell lung cancer | 119 (59/60) | 18682 | 500 (129/371) | Corrected |
| **GSE20189** | Non-small cell lung cancer | 162 (81/81) | 10949 | 500 (188/312) | Corrected |
| **GSE21933** | Non-small cell lung cancer | 42 (21/21) | 18941 | 500 (161/339) | Not detected |

| | | | | | |
|---|---|---|---|---|---|
| **GSE27262** | Non-small cell lung cancer | 50 (25/25) | 18369 | 500 (100/400) | Corrected |
| **GSE52248*** | Non-small cell lung cancer | 18 (12/6) | 15210 | 500 (153/347) | Not detected |
| **GSE16515** | Pancreatic cancer | 52 (36/16) | 18361 | 500 (374/126) | Corrected |
| **GSE18670** | Pancreatic cancer | 23 (11/12) | 18656 | 320 (187/133) | Corrected |
| **GSE23397** | Pancreatic cancer | 21 (15/6) | 15168 | 500 (218/282) | Corrected |
| **GSE28735** | Pancreatic cancer | 90 (45/45) | 18141 | 500 (378/122) | Not detected |
| **GSE42952** | Pancreatic cancer | 23 (11/12) | 18655 | 500 (131/369) | Not detected |
| **GSE18838** | Parkinson disease | 28 (17/11) | 14894 | 500 (134/366) | Corrected |
| **GSE19587†** | Parkinson disease | 22 (12/10) | 11219 | 500 (164/336) | Corrected |
| **GSE20141** | Parkinson disease | 18 (10/8) | 17802 | 500 (452/48) | Corrected |
| **GSE20146†** | Parkinson disease | 19 (10/9) | 17870 | 132 (25/107) | Corrected |
| **GSE20163** | Parkinson disease | 17 (8/9) | 11335 | 500 (177/323) | Corrected |
| **GSE20164†** | Parkinson disease | 11 (6/5) | 11239 | 195 (64/131) | Corrected |
| **GSE20291** | Parkinson disease | 35 (15/20) | 11341 | 500 (227/273) | Corrected |
| **GSE20292** | Parkinson disease | 29 (11/18) | 11335 | 500 (232/268) | Corrected |
| **GSE20314†** | Parkinson disease | 8 (4/4) | 11118 | 58 (38/20) | Corrected |
| **GSE20333†** | Parkinson disease | 12 (6/6) | 6756 | 298 (226/72) | Corrected |
| **GSE7621** | Parkinson disease | 25 (16/9) | 18430 | 500 (204/296) | Corrected |
| **GSE90514*** | Parkinson disease | 8 (4/4) | 14486 | 128 (84/44) | Not detected |
| **GSE11682†** | Prostate cancer | 33 (17/16) | 19547 | 59 (28/31) | Corrected |
| **GSE22260*** | Prostate cancer | 30 (20/10) | 15837 | 500 (329/171) | Not detected |
| **GSE30521** | Prostate cancer | 22 (17/5) | 15150 | 500 (292/208) | Corrected |
| **GSE10927** | Renal cell carcinoma | 64 (54/10) | 18392 | 500 (94/406) | Corrected |
| **GSE15641** | Renal cell carcinoma | 92 (69/23) | 11227 | 500 (103/397) | Corrected |
| **GSE33371** | Renal cell carcinoma | 64 (54/10) | 18392 | 500 (94/406) | Corrected |
| **GSE53757** | Renal cell carcinoma | 143 (71/72) | 18268 | 500 (173/327) | Corrected |

| | | | | | |
|---|---|---|---|---|---|
| **GSE55235** | Rheumatoid arthritis | 20 (10/10) | 11131 | 500 (320/180) | Corrected |
| **GSE30153** | Systemic lupus erythematosus | 26 (17/9) | 18065 | 20 (5/15) | Corrected |
| **GSE50635** | Systemic lupus erythematosus | 48 (32/16) | 17642 | 293 (214/79) | Corrected |
| **GSE48850*** | Thyroid cancer | 11 (6/5) | 15665 | 500 (249/251) | Not detected |
| **GSE10586†** | Type I diabetes mellitus | 27 (12/15) | 18023 | 25 (18/7) | Corrected |
| **GSE41762** | Type I diabetes mellitus | 77 (20/57) | 18094 | 500 (303/197) | Corrected |
| **GSE12643†** | Type II diabetes mellitus | 20 (10/10) | 7513 | 62 (49/13) | Corrected |
| **GSE13760†** | Type II diabetes mellitus | 21 (10/11) | 10994 | 29 (10/19) | Corrected |
| **GSE15653** | Type II diabetes mellitus | 18 (13/5) | 11157 | 500 (303/197) | Corrected |
| **GSE20966** | Type II diabetes mellitus | 20 (10/10) | 18819 | 240 (111/129) | Corrected |
| **GSE21340** | Type II diabetes mellitus | 20 (5/15) | 3499 | 444 (176/268) | Corrected |
| **GSE38642** | Type II diabetes mellitus | 63 (9/54) | 18105 | 500 (189/311) | Corrected |
| **GSE40234** | Type II diabetes mellitus | 62 (34/28) | 19098 | 452 (202/250) | Corrected |
| **GSE14858** | Acute myeloid leukemia | 39 (20/19) | 18337 | 500 (293/207) | Corrected |
| **GSE28619** | Alcoholic liver disease | 22 (15/7) | 18464 | 500 (331/169) | Not detected |
| **GSE12685** | Alzheimer disease | 13 (6/7) | 11141 | 500 (222/278) | Corrected |

**Supplementary Table 2 - Size of disease pathway networks at different levels of confidence.** We report the details of the disease/target pathways according to KEGG (*i.e.* pathway name, pathway id and pathway subclass), and the size of their network of interactions with all other pathways ("All", including the target pathway itself). As we assess the sensitivity of the methods by using the disease pathway network at different levels of confidence, we report the number of all linked pathways and for different top cutoffs (10,20,40). See "Disease pathway network construction" in the Methods section for details about the network construction process.

| Pathway name | Pathway ID | Node ID | Pathway subclass (Human diseases) | Top pathways | | | |
|---|---|---|---|---|---|---|---|
| | | | | **10** | **20** | **40** | **All** |
| **Acute myeloid leukemia** | hsa05221 | **5** | Cancer: specific types | 11 | 21 | 43 | 129 |
| **African trypanosomiasis** | hsa05143 | / | Infectious disease: parasitic | 11 | 21 | 42 | 73 |
| **AGE-RAGE signaling pathway in diabetic complications** | hsa04933 | / | Endocrine and metabolic disease | 12 | 21 | 42 | 143 |
| **Alcoholic liver disease** | hsa04936 | **17** | Endocrine and metabolic disease | 11 | 21 | 41 | 64 |
| **Alcoholism** | hsa05034 | / | Substance dependence | 11 | 21 | 41 | 106 |
| **Allograft rejection** | hsa05330 | / | Immune disease | 11 | 21 | 41 | 53 |
| **Alzheimer disease** | hsa05010 | **24** | Neurodegenerative disease | 11 | 21 | 41 | 82 |
| **Amoebiasis** | hsa05146 | / | Infectious disease: parasitic | 11 | 21 | 41 | 98 |
| **Amphetamine addiction** | hsa05031 | / | Substance dependence | 11 | 21 | 41 | 75 |
| **Amyotrophic lateral sclerosis** | hsa05014 | / | Neurodegenerative disease | 11 | 21 | 24 | 24 |
| **Arrhythmogenic right ventricular cardiomyopathy** | hsa05412 | / | Cardiovascular disease | 11 | 21 | 27 | 27 |
| **Asthma** | hsa05310 | **19** | Immune disease | 11 | 21 | 41 | 41 |
| **Autoimmune thyroid disease** | hsa05320 | / | Immune disease | 11 | 22 | 41 | 52 |
| **Bacterial invasion of epithelial cells** | hsa05100 | / | Infectious disease: bacterial | 11 | 21 | 41 | 80 |
| **Basal cell carcinoma** | hsa05217 | / | Cancer: specific types | 11 | 21 | 38 | 38 |
| **Bladder cancer** | hsa05219 | **9** | Cancer: specific types | 11 | 21 | 41 | 108 |
| **Breast cancer** | hsa05224 | **2** | Cancer: specific types | 11 | 21 | 42 | 109 |
| **Central carbon metabolism in cancer** | hsa05230 | / | Cancer: overview | 11 | 21 | 41 | 95 |
| **Chagas disease** | hsa05142 | / | Infectious disease: parasitic | 11 | 22 | 41 | 139 |

| | | | | | | | |
|---|---|---|---|---|---|---|---|
| **Chemical carcinogenesis - DNA adducts** | hsa05204 | **/** | Cancer: overview | 11 | 15 | 15 | 15 |
| **Chemical carcinogenesis - reactive oxygen species** | hsa05208 | **/** | Cancer: overview | 11 | 21 | 43 | 73 |
| **Chemical carcinogenesis - receptor activation** | hsa05207 | **/** | Cancer: overview | 12 | 22 | 41 | 106 |
| **Choline metabolism in cancer** | hsa05231 | **/** | Cancer: overview | 11 | 21 | 41 | 129 |
| **Chronic myeloid leukemia** | hsa05220 | **12** | Cancer: specific types | 11 | 22 | 41 | 129 |
| **Cocaine addiction** | hsa05030 | **/** | Substance dependence | 11 | 21 | 41 | 82 |
| **Colorectal cancer** | hsa05210 | **1** | Cancer: specific types | 11 | 21 | 41 | 128 |
| **Coronavirus disease - COVID-19** | hsa05171 | **/** | Infectious disease: viral | 11 | 21 | 41 | 98 |
| **Cushing syndrome** | hsa04934 | **/** | Endocrine and metabolic disease | 11 | 21 | 41 | 92 |
| **Diabetic cardiomyopathy** | hsa05415 | **/** | Cardiovascular disease | 12 | 21 | 41 | 50 |
| **Dilated cardiomyopathy** | hsa05414 | **14** | Cardiovascular disease | 11 | 21 | 41 | 66 |
| **Endometrial cancer** | hsa05213 | **10** | Cancer: specific types | 11 | 21 | 42 | 122 |
| **Epithelial cell signaling in Helicobacter pylori infection** | hsa05120 | **/** | Infectious disease: bacterial | 12 | 22 | 42 | 105 |
| **Epstein-Barr virus infection** | hsa05169 | **/** | Infectious disease: viral | 11 | 21 | 41 | 109 |
| **Fluid shear stress and atherosclerosis** | hsa05418 | **/** | Cardiovascular disease | 11 | 21 | 41 | 120 |
| **Gastric cancer** | hsa05226 | **/** | Cancer: specific types | 11 | 21 | 41 | 101 |
| **Glioma** | hsa05214 | **/** | Cancer: specific types | 11 | 21 | 41 | 129 |
| **Graft-versus-host disease** | hsa05332 | **/** | Immune disease | 12 | 21 | 41 | 53 |
| **Hepatitis B** | hsa05161 | **/** | Infectious disease: viral | 11 | 21 | 41 | 124 |
| **Hepatitis C** | hsa05160 | **22** | Infectious disease: viral | 12 | 21 | 41 | 128 |
| **Hepatocellular carcinoma** | hsa05225 | **11** | Cancer: specific types | 11 | 21 | 41 | 110 |
| **Herpes simplex virus 1 infection** | hsa05168 | **/** | Infectious disease: viral | 11 | 21 | 41 | 97 |
| **Human cytomegalovirus infection** | hsa05163 | **/** | Infectious disease: viral | 12 | 21 | 41 | 147 |
| **Human immunodeficiency virus 1 infection** | hsa05170 | **/** | Infectious disease: viral | 11 | 21 | 41 | 120 |
| **Human papillomavirus infection** | hsa05165 | **/** | Infectious disease: viral | 11 | 23 | 41 | 120 |
| **Human T-cell leukemia virus 1 infection** | hsa05166 | **/** | Infectious disease: viral | 11 | 21 | 41 | 114 |

| | | | | | | | |
|---|---|---|---|---|---|---|---|
| **Huntington disease** | hsa05016 | **26** | Neurodegenerative disease | 11 | 21 | 23 | 23 |
| **Hypertrophic cardiomyopathy** | hsa05410 | **/** | Cardiovascular disease | 11 | 21 | 41 | 43 |
| **Inflammatory bowel disease** | hsa05321 | **18** | Immune disease | 11 | 22 | 41 | 79 |
| **Influenza A** | hsa05164 | **/** | Infectious disease: viral | 11 | 21 | 41 | 105 |
| **Insulin resistance** | hsa04931 | **/** | Endocrine and metabolic disease | 11 | 21 | 41 | 119 |
| **Kaposi sarcoma-associated herpesvirus infection** | hsa05167 | **/** | Infectious disease: viral | 11 | 21 | 41 | 132 |
| **Legionellosis** | hsa05134 | **/** | Infectious disease: bacterial | 11 | 21 | 41 | 62 |
| **Leishmaniasis** | hsa05140 | **/** | Infectious disease: parasitic | 11 | 22 | 41 | 99 |
| **Lipid and atherosclerosis** | hsa05417 | **/** | Cardiovascular disease | 13 | 23 | 42 | 121 |
| **Malaria** | hsa05144 | **/** | Infectious disease: parasitic | 11 | 21 | 41 | 58 |
| **Maturity onset diabetes of the young** | hsa04950 | **/** | Endocrine and metabolic disease | 3 | 3 | 3 | 3 |
| **Measles** | hsa05162 | **23** | Infectious disease: viral | 11 | 21 | 41 | 113 |
| **Melanoma** | hsa05218 | **6** | Cancer: specific types | 11 | 21 | 41 | 114 |
| **MicroRNAs in cancer** | hsa05206 | **/** | Cancer: overview | 11 | 22 | 41 | 86 |
| **Morphine addiction** | hsa05032 | **/** | Substance dependence | 11 | 21 | 41 | 60 |
| **Nicotine addiction** | hsa05033 | **/** | Substance dependence | 11 | 21 | 23 | 23 |
| **Non-alcoholic fatty liver disease** | hsa04932 | **/** | Endocrine and metabolic disease | 11 | 21 | 41 | 96 |
| **Non-small cell lung cancer** | hsa05223 | **8** | Cancer: specific types | 12 | 21 | 41 | 129 |
| **Pancreatic cancer** | hsa05212 | **7** | Cancer: specific types | 11 | 21 | 41 | 135 |
| **Parkinson disease** | hsa05012 | **25** | Neurodegenerative disease | 11 | 21 | 28 | 28 |
| **Pathogenic Escherichia coli infection** | hsa05130 | **/** | Infectious disease: bacterial | 12 | 21 | 42 | 105 |
| **Pathways in cancer** | hsa05200 | **/** | Cancer: overview | 11 | 21 | 41 | 73 |
| **Pathways of neurodegeneration - multiple diseases** | hsa05022 | **/** | Neurodegenerative disease | 11 | 21 | 41 | 52 |
| **PD-L1 expression and PD-1 checkpoint pathway in cancer** | hsa05235 | **/** | Cancer: overview | 11 | 21 | 42 | 131 |
| **Pertussis** | hsa05133 | **/** | Infectious disease: bacterial | 11 | 21 | 42 | 108 |
| **Primary immunodeficiency** | hsa05340 | **/** | Immune disease | 11 | 21 | 26 | 26 |

| | | | | | | | |
|---|---|---|---|---|---|---|---|
| **Prion disease** | hsa05020 | **/** | Neurodegenerative disease | 11 | 21 | 41 | 77 |
| **Prostate cancer** | hsa05215 | **4** | Cancer: specific types | 11 | 22 | 42 | 115 |
| **Proteoglycans in cancer** | hsa05205 | **/** | Cancer: overview | 11 | 21 | 41 | 121 |
| **Renal cell carcinoma** | hsa05211 | **3** | Cancer: specific types | 11 | 21 | 41 | 129 |
| **Rheumatoid arthritis** | hsa05323 | **21** | Immune disease | 11 | 21 | 41 | 67 |
| **Salmonella infection** | hsa05132 | **/** | Infectious disease: bacterial | 11 | 21 | 41 | 114 |
| **Shigellosis** | hsa05131 | **/** | Infectious disease: bacterial | 11 | 22 | 41 | 114 |
| **Small cell lung cancer** | hsa05222 | **/** | Cancer: specific types | 11 | 21 | 41 | 109 |
| **Spinocerebellar ataxia** | hsa05017 | **/** | Neurodegenerative disease | 11 | 21 | 41 | 58 |
| **Staphylococcus aureus infection** | hsa05150 | **/** | Infectious disease: bacterial | 11 | 21 | 31 | 31 |
| **Systemic lupus erythematosus** | hsa05322 | **20** | Immune disease | 11 | 21 | 39 | 39 |
| **Thyroid cancer** | hsa05216 | **13** | Cancer: specific types | 11 | 21 | 41 | 108 |
| **Toxoplasmosis** | hsa05145 | **/** | Infectious disease: parasitic | 11 | 21 | 42 | 113 |
| **Transcriptional misregulation in cancer** | hsa05202 | **/** | Cancer: overview | 11 | 21 | 36 | 36 |
| **Tuberculosis** | hsa05152 | **/** | Infectious disease: bacterial | 11 | 21 | 41 | 108 |
| **Type I diabetes mellitus** | hsa04940 | **15** | Endocrine and metabolic disease | 11 | 21 | 41 | 43 |
| **Type II diabetes mellitus** | hsa04930 | **16** | Endocrine and metabolic disease | 11 | 21 | 41 | 125 |
| **Vibrio cholerae infection** | hsa05110 | **/** | Infectious disease: bacterial | 11 | 21 | 42 | 68 |
| **Viral carcinogenesis** | hsa05203 | **/** | Cancer: overview | 11 | 21 | 41 | 93 |
| **Viral myocarditis** | hsa05416 | **/** | Cardiovascular disease | 12 | 21 | 41 | 68 |
| **Yersinia infection** | hsa05135 | **/** | Infectious disease: bacterial | 11 | 21 | 42 | 124 |

**Supplementary Table 3 - Average disease pathway similarity within KEGG subclasses.** Listed is the average similarity, in terms of Jaccard index, between all disease pathways within the same KEGG subclass, for disease pathways in the Disease Pathway Network using the top 20 linked pathways.

| KEGG subclass | avg JI |
|---|---|
| **Cancer: specific types** | 0.45 |
| **Infectious disease: viral** | 0.40 |
| **Immune disease** | 0.40 |
| **Infectious disease: parasitic** | 0.33 |
| **Neurodegenerative disease** | 0.32 |
| **Substance dependence** | 0.28 |
| **Infectious disease: bacterial** | 0.20 |
| **Cardiovascular disease** | 0.18 |
| **Cancer: overview** | 0.16 |
| **Endocrine and metabolic disease** | 0.08 |

**Supplementary Table 4 - Summary of performances with top 20 links per target pathway in the disease pathway network.** True Positive (TP), False Negative (FN), False Positive (FP), True Negative (TN), True Positive Rate (TPR), False Positive Rate (FPR), G-mean and 1 - relative rank are extracted over all predictions.

| Method | TP | FN | FP | TN | 1-Rank | TPR | FPR | G-mean |
|---|---|---|---|---|---|---|---|---|
| **ANUBIX-FunCoup** | 611 | 1118 | 3526 | 48344 | 0.60 | 0.35 | 0.07 | 0.57 |
| **ANUBIX-STRING** | 610 | 1119 | 3720 | 48150 | 0.62 | 0.35 | 0.07 | 0.57 |
| **BinoX-FunCoup** | 811 | 918 | 10546 | 41324 | 0.61 | 0.47 | 0.20 | 0.61 |
| **BinoX-STRING** | 977 | 752 | 16665 | 35205 | 0.65 | **0.57** | **0.32** | **0.62** |
| **CAMERA** | 489 | 1240 | 463 | 51407 | 0.48 | 0.28 | 0.01 | 0.53 |
| **CAMERA*** | 177 | 1552 | 662 | 51208 | 0.51 | 0.10 | 0.01 | 0.32 |
| **CePa-ORA** | 349 | 1380 | 4971 | 46899 | 0.60 | 0.20 | 0.10 | 0.43 |
| **EASE** | 145 | 1584 | 525 | 51345 | 0.57 | 0.08 | 0.01 | 0.29 |
| **fGSEA** | 684 | 1045 | 2616 | 49254 | 0.52 | 0.40 | 0.05 | 0.61 |
| **Fisher** | 190 | 1539 | 1584 | 50286 | 0.53 | 0.11 | 0.03 | 0.33 |
| **GSA** | 187 | 1542 | 2179 | 49691 | 0.53 | 0.11 | 0.04 | 0.32 |
| **GSEA** | 684 | 1045 | 2631 | 49239 | 0.53 | 0.40 | 0.05 | 0.61 |
| **GSVA** | 680 | 1049 | 13104 | 38766 | 0.51 | 0.39 | 0.25 | 0.54 |
| **NEAT-FunCoup** | 775 | 954 | 10046 | 41824 | 0.60 | 0.45 | 0.19 | 0.60 |
| **NEAT-STRING** | 950 | 779 | 15241 | 36629 | 0.64 | 0.55 | 0.29 | **0.62** |
| **netPEA-FunCoup** | 560 | 1169 | 6320 | 45550 | 0.58 | 0.32 | 0.12 | 0.53 |
| **netPEA-STRING** | 621 | 1108 | 5635 | 46235 | 0.65 | 0.36 | 0.11 | 0.57 |
| **PADOG** | 188 | 1541 | 3051 | 48819 | 0.57 | 0.11 | 0.06 | 0.32 |
| **ROAST** | 738 | 991 | 18499 | 33371 | 0.51 | 0.43 | 0.36 | 0.52 |
| **SPIA** | 206 | 1523 | 1709 | 50161 | **0.68** | 0.12 | 0.03 | 0.34 |

**Supplementary Table 5 - Spearman correlation *p*-values between pathway network properties and false positive rate.** The correlation *p*-values were estimated with a test for correlation between paired samples for a selection of network properties against the False Positive Rate (FPR) of KEGG pathways as predicted by network-based EA methods with FunCoup and STRING as underlying functional association networks. Significant *p*-values below the cutoff 0.05 are marked in bold. See Supplementary Figure 5 to find the respective Spearman correlations.

| | FunCoup | | | | STRING | | | |
|---|---|---|---|---|---|---|---|---|
| | **ANUBIX** | **BinoX** | **NEAT** | **netPEA** | **ANUBIX** | **BinoX** | **NEAT** | **netPEA** |
| **Density of hubs** | 9.6e-01 | **1.6e-12** | **8.3e-07** | **1.5e-04** | **1.5e-09** | **1.6e-35** | **1.0e-28** | **2.5e-06** |
| **Density of links** | 1.5e-01 | **5.8e-10** | **3.1e-07** | 1.2e-01 | **6.4e-08** | **1.3e-02** | 9.1e-02 | 2.1e-01 |
| **Fraction of hubs** | **3.6e-02** | **2.1e-45** | **2.3e-32** | **2.9e-07** | **6.1e-18** | **2.2e-74** | **9.5e-60** | **7.8e-07** |
| **Fraction of intralinks** | 2.8e-01 | **5.8e-41** | **1.7e-50** | 9.7e-01 | **4.6e-10** | **1.8e-03** | **3.1e-06** | **2.2e-04** |
| **Max degree** | 6.3e-01 | **7.2e-09** | **9.2e-05** | **3.8e-04** | **6.7e-30** | **3.7e-39** | **3.1e-29** | **2.8e-15** |
| **Mean clustering coefficient** | **2.5e-04** | **6.8e-04** | **1.4e-02** | 8.4e-01 | **1.2e-12** | **6.9e-07** | **3.0e-05** | **3.2e-05** |
| **Median degree** | 3.6e-01 | **3.2e-19** | **2.1e-13** | **2.3e-06** | **1.2e-10** | **2.4e-38** | **6.9e-31** | **4.3e-07** |
| **Number of CC** | 7.3e-01 | 8.6e-01 | 5.7e-01 | 3.9e-01 | 1.5e-01 | 9.8e-02 | 5.8e-02 | 5.2e-02 |
| **Size** | **1.7e-02** | **8.6e-36** | **1.3e-33** | **8.5e-04** | **2.1e-15** | **7.6e-55** | **4.1e-47** | **3.2e-04** |

### References


1. Wang JZ, Du Z, Payattakool R, et al. A new method to measure the semantic similarity of GO terms. Bioinformatics 2007; 23:1274–1281
2. Goeman JJ, Bühlmann P. Analyzing gene expression data in terms of gene sets: methodological issues. Bioinformatics 2007; 23:980–987
3. Krämer A, Green J, Pollard J Jr, et al. Causal analysis approaches in Ingenuity Pathway Analysis. Bioinformatics 2014; 30:523–530
4. Hosack DA, Dennis G Jr, Sherman BT, et al. Identifying biological themes within lists of genes with EASE. Genome Biol. 2003; 4:R70
5. Subramanian A, Tamayo P, Mootha VK, et al. Gene set enrichment analysis: a knowledge-based approach for interpreting genome-wide expression profiles. Proc. Natl. Acad. Sci. U. S. A. 2005; 102:15545–15550
6. Korotkevich G, Sukhov V, Budin N, et al. Fast gene set enrichment analysis. bioRxiv 2021;
7. Efron B, Tibshirani R. On testing the significance of sets of genes. Ann. Appl. Stat. 2007; 1:107–129
8. Wu D, Lim E, Vaillant F, et al. ROAST: rotation gene set tests for complex microarray experiments. Bioinformatics 2010; 26:2176–2182
9. Hänzelmann S, Castelo R, Guinney J. GSVA: gene set variation analysis for microarray and RNA-seq data. BMC Bioinformatics 2013; 14:7
10. Wu D, Smyth GK. Camera: a competitive gene set test accounting for inter-gene correlation. Nucleic Acids Res. 2012; 40:e133
11. Tarca AL, Draghici S, Bhatti G, et al. Down-weighting overlapping genes improves gene set analysis. BMC Bioinformatics 2012; 13:136
12. Tarca AL, Draghici S, Khatri P, et al. A novel signaling pathway impact analysis. Bioinformatics 2009; 25:75–82
13. Gu Z, Wang J. CePa: an R package for finding significant pathways weighted by multiple network centralities. Bioinformatics 2013; 29:658–660
14. Nguyen T-M, Shafi A, Nguyen T, et al. Identifying significantly impacted pathways: a comprehensive review and assessment. Genome Biol. 2019; 20:1–15
15. Liu L, Ruan J. Network-based Pathway Enrichment Analysis. Proceedings 2013; 218–221
16. Signorelli M, Vinciotti V, Wit EC. NEAT: an efficient network enrichment analysis test. BMC Bioinformatics 2016; 17:352
17. Ogris C, Guala D, Helleday T, et al. A novel method for crosstalk analysis of biological networks: improving accuracy of pathway annotation. Nucleic Acids Res. 2017; 45:e8
18. Castresana-Aguirre M, Sonnhammer ELL. Pathway-specific model estimation for improved pathway annotation by network crosstalk. Sci. Rep. 2020; 10:13585
